# The Amphibian Genomics Consortium: advancing genomic and genetic resources for amphibian research and conservation

**DOI:** 10.1101/2024.06.27.601086

**Authors:** Tiffany A. Kosch, María Torres-Sánchez, H. Christoph Liedtke, Kyle Summers, Maximina H. Yun, Andrew J. Crawford, Simon T. Maddock, Md. Sabbir Ahammed, Victor L. N. Araújo, Lorenzo V. Bertola, Gary M. Bucciarelli, Albert Carné, Céline M. Carneiro, Kin O. Chan, Ying Chen, Angelica Crottini, Jessica M. da Silva, Robert D. Denton, Carolin Dittrich, Gonçalo Espregueira Themudo, Katherine A. Farquharson, Natalie J. Forsdick, Edward Gilbert, Jing Che, Barbara A. Katzenback, Ramachandran Kotharambath, Nicholas A. Levis, Roberto Márquez, Glib Mazepa, Kevin P. Mulder, Hendrik Müller, Mary J. O’Connell, Pablo Orozco-terWengel, Gemma Palomar, Alice Petzold, David W. Pfennig, Karin S. Pfennig, Michael S. Reichert, Jacques Robert, Mark D. Scherz, Karen Siu-Ting, Anthony A. Snead, Matthias Stöck, Adam M. M. Stuckert, Jennifer L. Stynoski, Rebecca D. Tarvin, Katharina C. Wollenberg Valero, The Amphibian Genomics Consortium (AGC)

## Abstract

Amphibians represent a diverse group of tetrapods, marked by deep divergence times between their three systematic orders and families. Studying amphibian biology through the genomics lens increases our understanding of the features of this animal class and that of other terrestrial vertebrates. The need for amphibian genomic resources is more urgent than ever due to the increasing threats to this group. Amphibians are one of the most imperiled taxonomic groups, with approximately 41% of species threatened with extinction due to habitat loss, changes in land use patterns, disease, climate change, and their synergistic effects. Amphibian genomic resources have provided a better understanding of ontogenetic diversity, tissue regeneration, diverse life history and reproductive modes, anti-predator strategies, and resilience and adaptive responses. They also serve as essential models for studying broad genomic traits, such as evolutionary genome expansions and contractions, as they exhibit the widest range of genome sizes among all animal taxa and possess multiple mechanisms of genetic sex determination. Despite these features, genome sequencing of amphibians has significantly lagged behind that of other vertebrates, primarily due to the challenges of assembling their large, repeat-rich genomes and the relative lack of societal support. The emergence of long-read sequencing technologies, combined with advanced molecular and computational techniques that improve scaffolding and reduce computational workloads, is now making it possible to address some of these challenges. To promote and accelerate the production and use of amphibian genomics research through international coordination and collaboration, we launched the Amphibian Genomics Consortium (AGC, https://mvs.unimelb.edu.au/amphibian-genomics-consortium) in early 2023. This burgeoning community already has more than 282 members from 41 countries. The AGC aims to leverage the diverse capabilities of its members to advance genomic resources for amphibians and bridge the implementation gap between biologists, bioinformaticians, and conservation practitioners. Here we evaluate the state of the field of amphibian genomics, highlight previous studies, present challenges to overcome, and call on the research and conservation communities to unite as part of the AGC to enable amphibian genomics research to “leap” to the next level.

## State of the field of amphibian genomics

In 2010, the genome of the Western clawed frog (*Xenopus tropicalis*) was sequenced, marking the first genome assembly for Class Amphibia [1]. This species serves as a crucial laboratory model organism for cell biology, molecular genetics, and developmental biology [2]. The first amphibian genome assembly came years after the completion of the first genomes for other vertebrate groups: fishes in 2002 (*Fugu rubripes*; [3]), mammals in 2003 (*Homo sapiens*; [4]), birds in 2004 (*Gallus gallus*; [5]), and reptiles in 2007 (*Anolis carolinensis*; Anolis Genome Project https://www.broadinstitute.org/anolis/anolis-genome-project). Since then, the generation and annotation of amphibian reference genomes has dramatically lagged behind those of other vertebrates [6], even though amphibians represent nearly 22% of all tetrapods [7]. Nearly 15 years later, amphibians are still the tetrapod class with the lowest number of sequenced genomes (111 genomes of 8648 described amphibian species being the tetrapod class with the second lowest proportion after non-avian Reptiles, i.e. crocodilians, lepidosaurs, and testudines [database records accessed on 1 March 2024], Fig. 1A and Supplementary File 1). This is likely attributable to the size of amphibian genomes, which are generally larger than the genomes of other terrestrial vertebrates (Fig. 1B and Fig. S1; see Supplementary Material for methodological information). Indeed, among all vertebrates, only the genomes of lungfish are larger (up to 130 Gb) than the largest amphibian genomes (up to ∼120 Gb in *Necturus lewisi*) [8–11].

**Figure 1.**
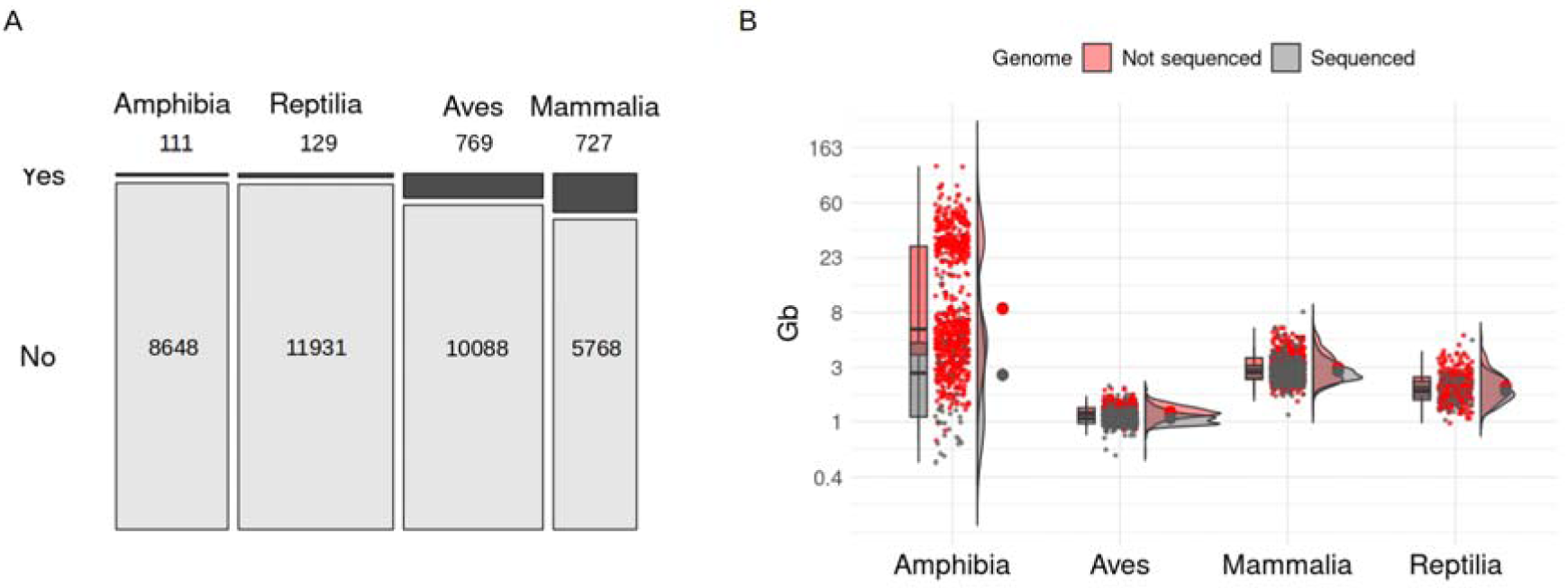
Estimated genome size across tetrapod classes in relation to sequenced genomes. (A) Mosaic plot representing the percentage of species with sequenced genomes as a proportion of the number of described species for each tetrapod class (Yes: % species with sequenced genome; No: % species without sequenced genome). (B) Combined box and density plot with raw data as points comparing genome size of species with sequenced genome (gray; genome sizes from NCBI genome assemblies) versus a subset of species without a sequenced genome (red; genome sizes from the Animal Genome Size Database) for each tetrapod class. The y-axis is log-transformed to facilitate visualization. Information about sequenced genomes and genome sizes was obtained from the NCBI Genome Browser, the Animal Genome Size database, and amphibian records from [12, 20].

To reduce costs and enhance feasibility, early amphibian genome sequencing projects tended to select species with comparatively small genomes (Fig. 1B). This has resulted in disproportionately fewer sequenced salamander genomes, given this is the amphibian order with the largest genomes [12]. To date, the largest amphibian genome assemblies belong to three salamander species: *Ambystoma mexicanum* (27.3 Gb assembly; [13]), *Pleurodeles waltl* (20.3 Gb; [14]), and *Calotriton arnoldi* (22.8 Gb; [15]). However, these only represent the lower end of the genome size range for this group, with the genomes of *Necturus* salamanders exceeding 100 Gb (Fig. 2) [10].

**Figure 2.**
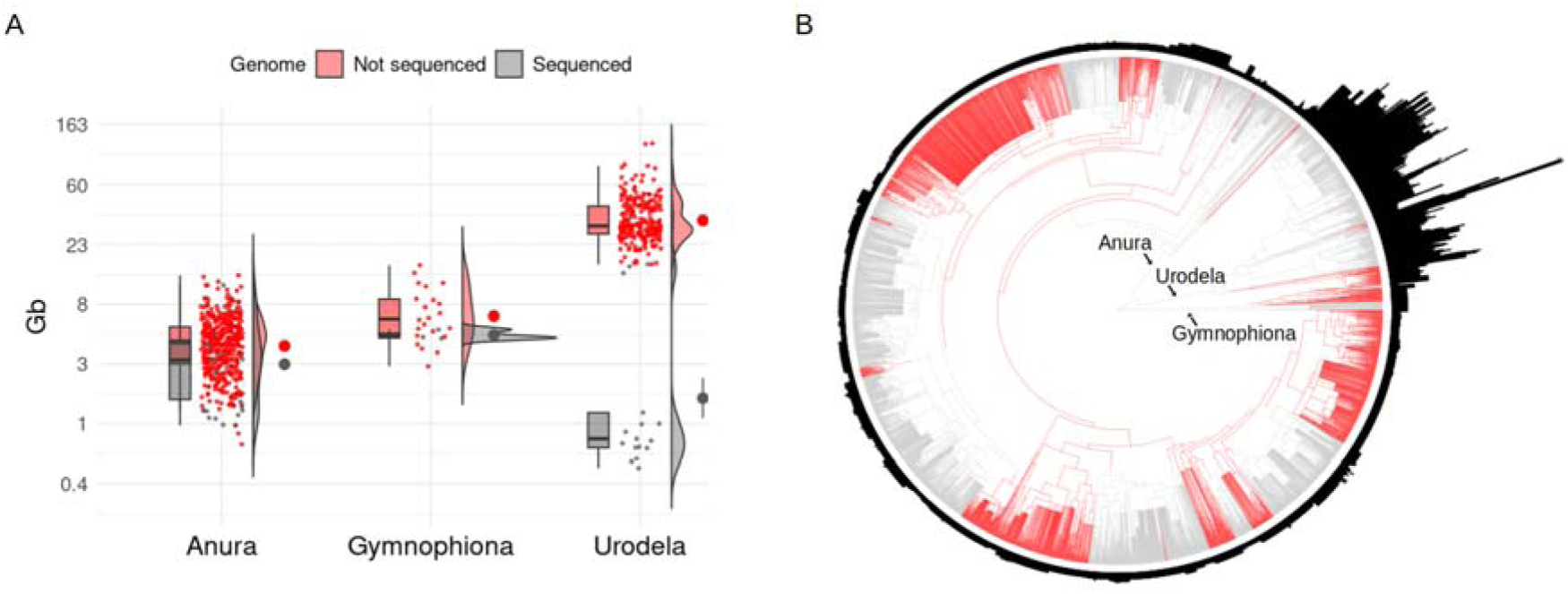
Estimated genome size across amphibian orders in relation to sequenced genomes. (A) Combined box and density plot with raw data as points showing genome size of species with sequenced genome (gray color; genome sizes from NCBI genome assemblies) versus a subset of species without available genome assembly (red color; genome sizes from the Animal Genome Size Database) for each amphibian order. The y-axis is logarithmic transformed to facilitate visualization. Information about sequenced genomes and genome sizes was obtained from the NCBI Genome Browser, the Animal Genome Size database [20], and amphibian records from [12]. (B) Amphibian phylogenetic tree was adapted from [71], which includes species with genome size estimates from Genomes on a Tree (GoaT) [19]. Branches are color coded to represent families without any genomic record (in red) and families with at least a representative genome sequenced (in gray). Bar plots around the phylogeny indicate relative genome sizes.

In addition to their large sizes, amphibian genomes have also been challenging to assemble due to their extensive repeat content (up to 82% [16]). Amphibian transposable elements have expanded and become highly abundant in younger clades, posing challenges for the construction of contiguous genome assemblies [17]. These characteristics of amphibian genomes make sequencing and assembly both costly and technically challenging (e.g., repetitive regions can often lead to fragmented assemblies when using short-read sequencing). However, the advent of new sequencing approaches such as long-read sequencing (e.g., PacBio HiFi and Oxford Nanopore Duplex), Hi-C scaffolding, along with reduced sequencing costs have resolved many of these assembly challenges (e.g., *Nanorana parkeri*; [18]).

Thus, the number of amphibian genome assemblies has increased rapidly in recent years, reaching 111 listed in early 2024 as reference genomes at the scaffold level or higher in the National Center for Biotechnology Information (NCBI) genome database (52 for Anura, 55 for Urodela, and four for Gymnophiona; NCBI genome database records accessed on 1 March 2024). Despite this rapid increase, the quality of available amphibian genomes varies significantly: only 38 are chromosome-level assemblies, and among these, only 16 are annotated. This indicates that the majority of available assemblies are incomplete or partial. For example, several recently published salamander genomes of the genus *Desmognathus* have assembly sizes of ∼1 Gb while their genome size estimates based on flow cytometry or image densitometry average 14 Gb [19, 20]. Furthermore, the gene content values for many of these incomplete genomes can be as low as 0.7% [16]. Besides the variation in quality, there are substantial taxonomic gaps in genome representation across Amphibia. Notably, 48 of the 77 amphibian families (62%) lack a representative genome assembly in the NCBI genome database (Fig. 2B), indicating significant gaps in our understanding (see “The AGC’s genome sequencing targets” section and Table 1 for more information about these 48 families).

**Table 1.**
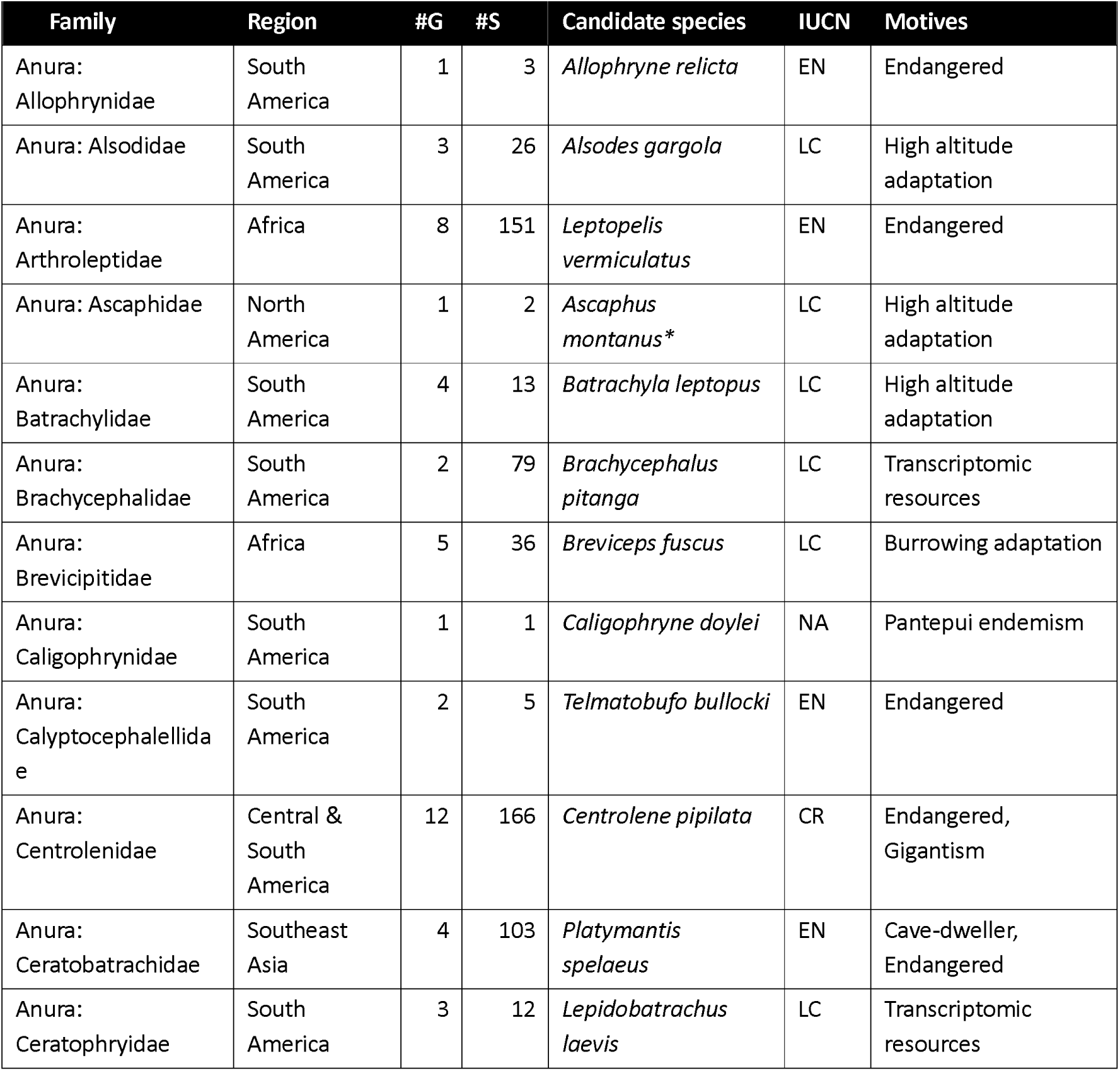
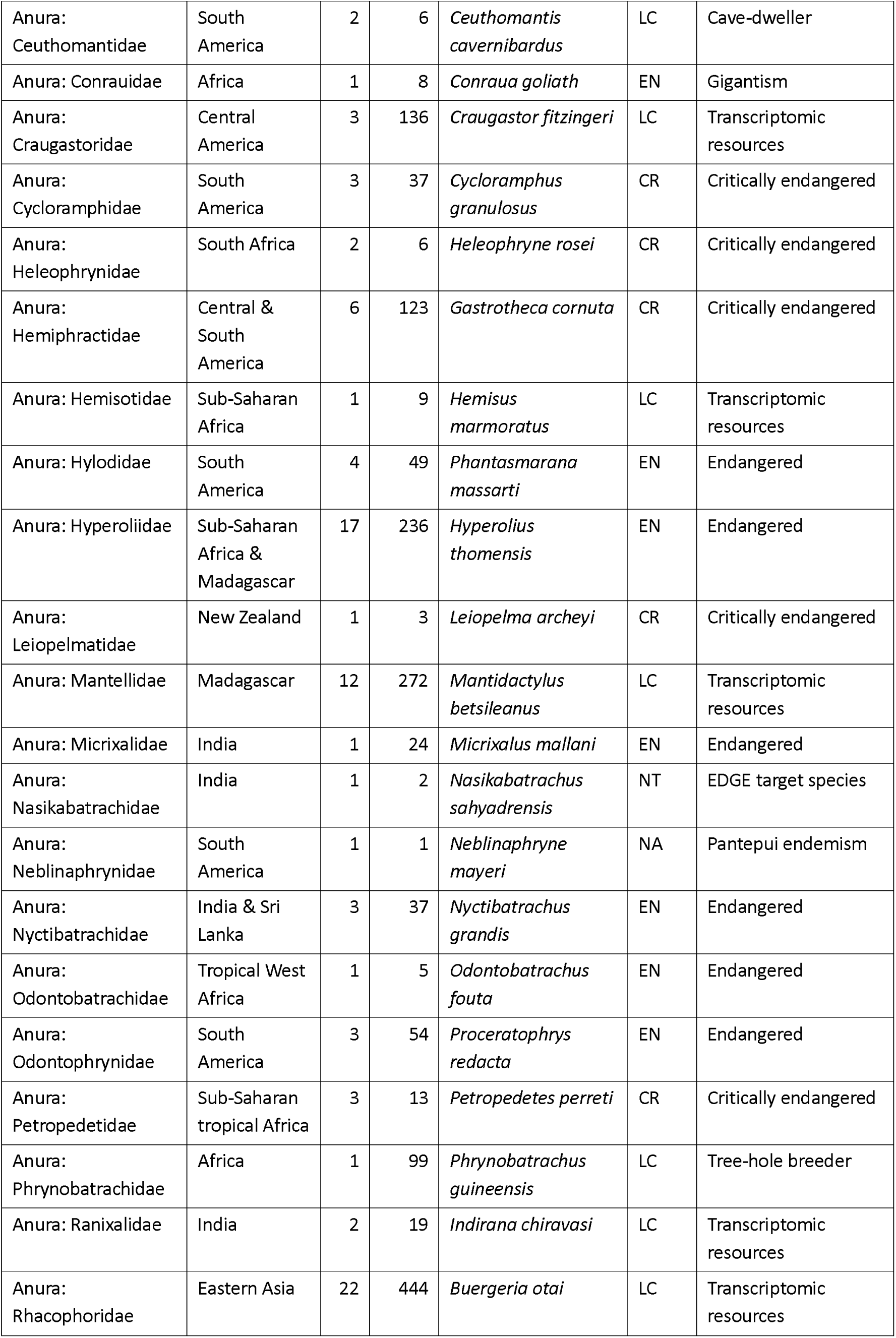
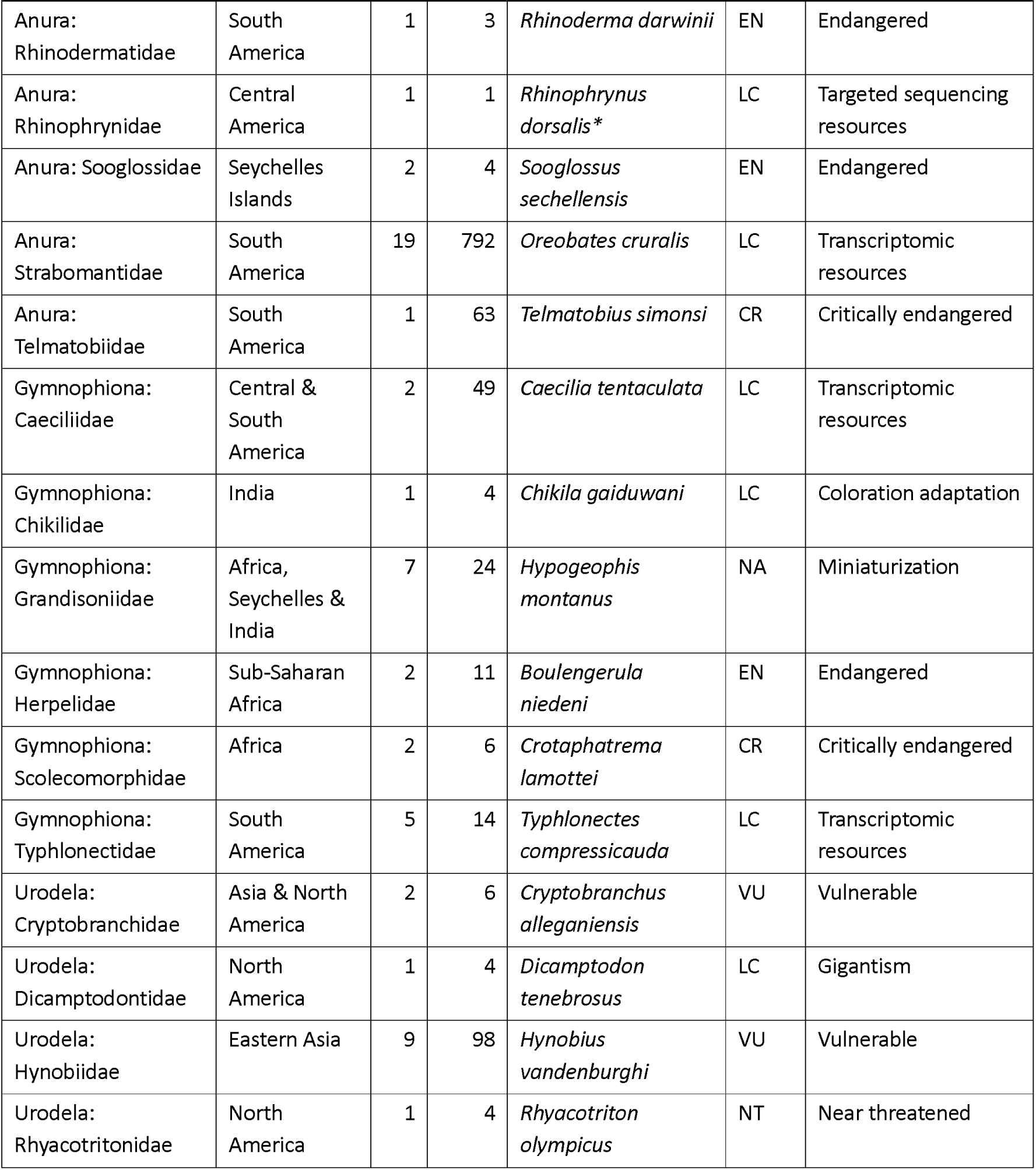
Amphibian Genomics Consortium (AGC) sequencing priority list. Table of amphibian families without any sequenced genomes. For each family, AGC proposed a candidate species based on its IUCN Red List category (LC: Least Concern, NT: Near Threatened, VU: Vulnerable, EN: Endangered, CR: Critically Endangered, and NA: Not evaluated), ecological and evolutionary distinctiveness, and availability of other genomic records. This table shows the amphibian order to which each family belongs and its number of genera (#G) and described extant species (#S) as well as distribution region. *Species with available draft genome assemblies in the GenomeArk (https://www.genomeark.org/).

Due to the difficulty of assembling genomes, most previous genomic research in amphibians has relied on alternative high-throughput sequencing methodologies, including RNA sequencing (RNA-seq), reduced representation or target-capture approaches, or metagenomic methods (Fig. 3 and Supplementary File 2 that contains the information for the search term “Amphibia” of the NCBI Sequence Read Archive [SRA] accessed on 1 March 2024). For example, RNA-Seq techniques have been used to explore gene expression across more than 300 different amphibian species (see Supplementary File 2 and Supplementary Methods for detail information about how SRA records were summarized). Furthermore, a substantial number of *de novo* transcriptomes are available through the NCBI Transcriptome Shotgun Assemblies (TSA) database (79 total: 59 for Anura, 15 for Urodela, and 5 for Gymnophiona). Various reduced-representation (e.g., ddRADseq) and targeted-capture sequencing approaches have also been implemented in recent years to obtain genome-wide sequence information from more than 1,400 amphibian species (see Supplementary File 2 and Supplementary Methods for detail information about how SRA records were summarized). All this information—from whole genomes to gene transcript features—has advanced the understanding of amphibian biology and directly contributed to conservation efforts as described below.

**Figure 3.**
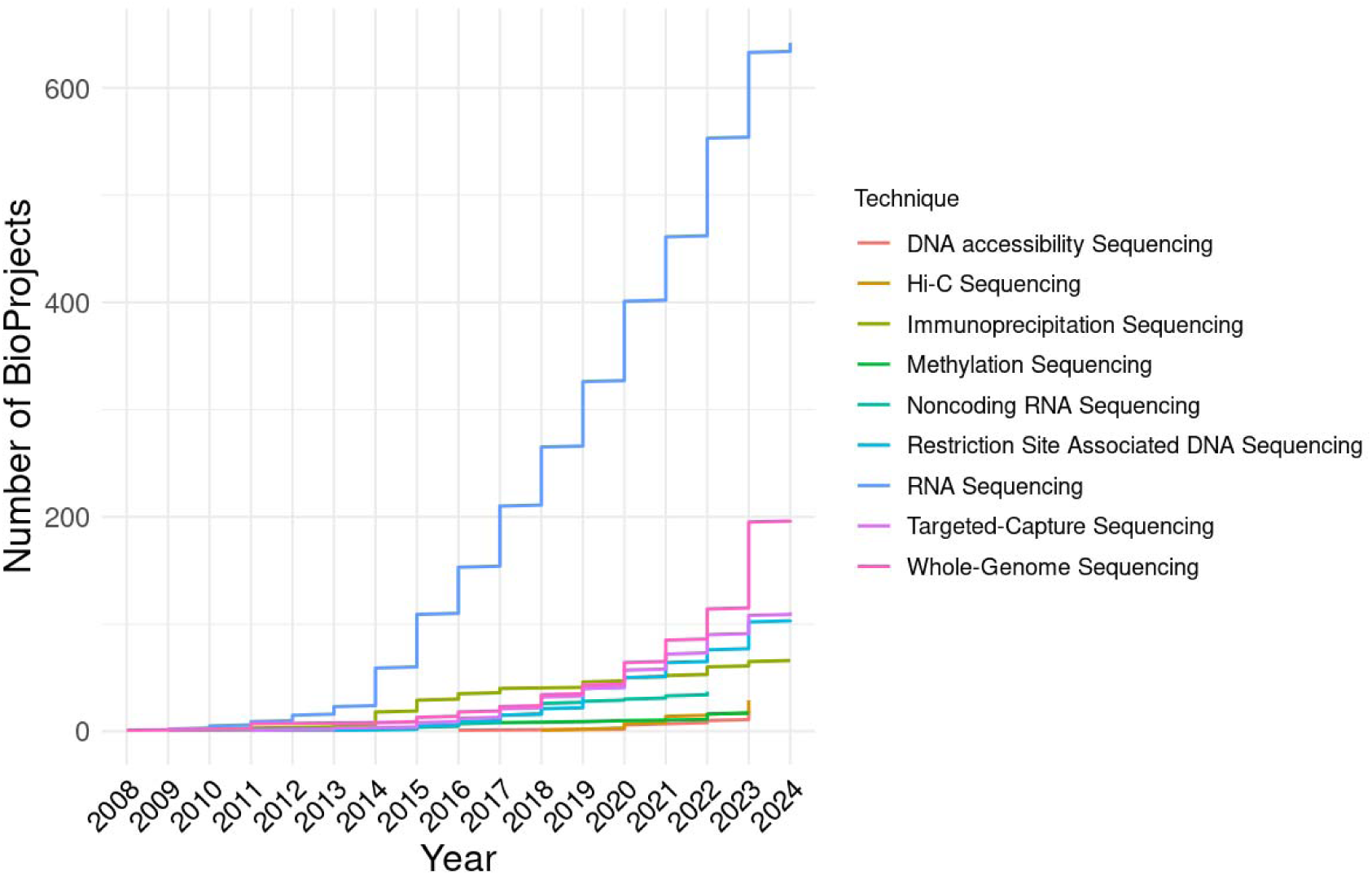
Main sequencing techniques applied to amphibian genomics studies. Yearly cumulative number of amphibian BioProjects split and color-coded by sequencing technique (DNA accessibility Sequencing includes ATAC-Seq and Mnase-Seq; Immunoprecipitation Sequencing includes: ChIP-Seq and RIP-Seq; Amplicon sequencing was included with Targeted-Capture Sequencing; Noncoding RNA Sequencing includes: miRNA-Seq and ncRNA-Seq). BioProject information was obtained from the NCBI Sequence Read Archive (SRA, accessed 1 March 2024).

## Advancing research and conservation through amphibian genomics

Amphibians have many unique characteristics that make them subjects of interest to a wide variety of scientific disciplines, spanning from developmental biology and medical research to ecology and evolution. The rapid development of genomic tools is galvanizing the study of amphibian biology and uncovering important facets of their biology and conservation [21–23]. We highlight some examples here and state the imperious need to generate amphibian genomic resources to decrease further biodiversity loss as the ultimate reason.

### Embryogenesis, developmental and regenerative biology

Amphibians have played a fundamental role in uncovering developmental principles [for a detailed review see 24]. Research on anurans has enabled the understanding of critical developmental mechanisms such as the breaking of egg asymmetry [25], axis establishment, and nerve transmission [26]. Notably, the availability of genome assemblies for *Xenopus laevis* and *X. tropicalis* has significantly advanced embryological and developmental biology. This advancement has enabled gene loss-of-function research through the combination of transgenesis with RNA interference, gene editing, and enhanced morpholino design. This has facilitated the in-depth analysis of regulatory and non-coding genomic influences in developmental processes [27, 28]. Consequently, these studies have generated thousands of genomic and transcriptomic resources for these two species [29, 30].

Yet, there is much more to uncover about amphibian development, especially given the numerous developmental modalities found across amphibians, which likely demonstrates the highest diversity among vertebrates [31]. This includes direct development (egg to froglet; the first genome of a direct-developing amphibian, *Eleutherodactylus coqui*, was published in 2024 [32]), and phenotypic plasticity [33, 34].

Sexual development and determination are also diverse and unique in amphibians [35]. Unlike most mammals and birds who have degenerate Y and W chromosomes, most amphibians have undifferentiated sex chromosomes, making it extremely difficult to study sex evolution through traditional cytogenetic techniques [36, 37]. However, sex-determining systems are starting to be explored through high-throughput sequencing [6, 38–42]. For example, the application of multiple omics techniques led to the identification of a Y-specific non-coding RNA in the 5’-region of the *bod1l* gene, which is involved in male sex determination in *Bufotes viridis* [41].

Strikingly, some salamanders in the genus *Ambystoma* exist as a single all-female, polyploid lineage that can incorporate new chromosome sets from up to five other sexual species [43]. Transcriptomes from these salamanders have shown that gene expression from their divergent genomes is balanced for some genes but biased for others [44]. Sexual development in amphibians can result in sexually dimorphic features such as nuptial spines, which have been explored using comparative genomics approaches such as in the frog *Leptobrachium leishanense* [45].

The increasing availability of amphibian genomes will enable a deeper understanding of the molecular mechanisms underlying such ontogenetic diversity. Chromosome-level reference genomes provide high-resolution data crucial for identifying sex-determining regions, revealing new insights about these processes and, helping to address challenges of sex reversal due to temperature fluctuations and the increasing presence of endocrine disruptors [46].

Metamorphosis sets many amphibian species apart from amniotes. Transcriptomics has revealed a remarkable turnover in gene expression between larval and adult stages of both frogs [47–50] and salamanders [51, 52]. This represents genomic uncoupling of these life history phases with major macroevolutionary implications [49, 53]. Amphibian omics approaches are rapidly increasing our understanding of the developmental process of metamorphosis, including the role of methylation in gene regulation and other epigenetic markers [54]. Amphibians have also been found to respond to environmental perturbations by altering their behavior or phenotypes in various ways. These mechanisms, including change developmental rate [33], hybridization with positive fitness effects [55], production of novel trophic morphologies [56], and kin recognition to avoid cannibalizing relatives [57–59], remain poorly understood, and would benefit from further genomic research.

Due to their exceptional tissue repair and regenerative capacities [60, 61], amphibians are leading models for understanding the mechanisms of regeneration. This is particularly true for salamanders, which display the most extensive adult regenerative repertoire among vertebrates, including the ability to regenerate parts of their eyes, brain, heart, jaws, lungs, spinal cord, tail, and entire limbs [61]. Due to new genome assemblies for urodele species, *Ambystoma mexicanum* and *Pleurodeles waltl*, regeneration can now be studied with transgenesis, advanced imaging, and genome editing. Intensive transcriptomic sequencing for these two salamander species has facilitated gene expression studies, including investigations into regeneration processes and characterization of other genomic features [62]. Additionally, a novel mechanism of telomere length maintenance and elongation has recently been described in *P. waltl* [63] and, potentially linking regenerative capability with longevity. Other amphibian species have also contributed to genomic research on regeneration, for example, an early database compiled from gene expression resources of *Notophthalmus viridescens* [64].

### Ecology and evolution

Modern amphibians are the sister lineage of all amniotes, making them a valuable resource for studying species relationships and trait evolution. This is exemplified by studies that explore the rapid diversification of frogs [65], the evolution of vision [66], hybridogenesis [67–69], and the evolution of limblessness [70]. Amphibian phylogenomics has addressed many longstanding questions in amphibian evolution [71–74]. Comparative genomic analyses including amphibian groups have also revealed important gaps in our understanding of tetrapod molecular evolution such as chromosomal rearrangements and group-specific gene families that remain unclassified to date [70, 75, 76]. Nevertheless, there are numerous open questions and unresolved evolutionary relationships that could benefit from high-quality genomes, which are especially powerful in revealing the role of transposable elements in adaptation and evolution [77]. In this section, we explore how genomics is being applied to understand the diverse ecological and evolutionary features unique to amphibians.

Like mammals, birds, and reptiles [78–80], some amphibians have evolved the ability to live in high-elevation environments such as the Andes (up to 5400 m) [81, 82] and the Tibetan Plateau (4478 m) [18]. However, unlike other groups, amphibians lack fur, feathers, or scales to protect them from physiological stressors such as UV exposure. This vulnerability makes them an intriguing model for studying the effects of UV radiation, which is relevant not only to humans [18] but also to species impacted by climate change. Amphibians have evolved multiple mechanisms of resisting UV, including increasing antioxidant efficiency and gene regulatory changes in defense pathways [18, 83]. There is evidence that genes that impact other high-elevation traits (e.g., hypoxia resistance, immunity, cold tolerance) have evolved convergently across distantly related families (e.g., Dicroglossidae, Bufonidae, Megophryidae, Ranidae) [84, 85], and that intraspecific divergence in many of these genes correlates with elevation deepening our understanding of evolutionary processes shaped by environmental conditions [86, 87]. While we are beginning to understand the genetic mechanisms of high-elevation adaptation in some Asian and North American frogs, this has yet to be investigated in other high-elevation amphibians where genomic data is still missing, including Andean anurans (e.g., *Telmatobius culeus* [88]) and high-elevation salamanders, such as *Pseudoeurycea gadovii* [89]).

The ability to produce or sequester toxins has evolved across all three amphibian orders, where it primarily serves as an anti-predation mechanism. The source of amphibian toxins varies: some species are capable of synthesizing poisonous compounds (e.g., bufonids, myobatrachids), whereas others sequester toxic substances from their diet (e.g., dendrobatids, mantellids) [90–93] or microbial symbionts (e.g., newts) [94]. Since dendrobatid frogs sequester their toxins from prey (e.g., mites and ants), they lack genes encoding these toxins [95, 96]. However, they require genes to facilitate the transport of these toxins to the skin. Recent genomic and proteomic research has identified candidate genes coding for proteins that may serve dual roles in toxin transport and resistance [97–99]. Comparative genomic research has identified specific substitutions that allow toxic amphibian species to effectively mitigate the effects of the sequestered toxins on their own tissues [100–102]. Skin transcriptomes have also proven to be a rich source for data mining and the identification of candidate toxins and antimicrobial peptides in various amphibians [103–107], which could potentially be used for future human medical treatments.

Interactions between toxic amphibians and their predators have resulted in a fascinating variety of co-evolutionary arms races. These include well-characterized systems of toxicity resistance mechanisms in amphibian predators [108–112] and aposematism and mimicry in toxic species [113, 114]. Research on aposematism and mimicry has utilized whole genome, exome capture, and transcriptome sequencing to elucidate the genes underlying the vast diversity of color patterns across populations and species in dendrobatids [115–120]. These approaches have yielded a goldmine of information that can be used to understand the genes, gene networks, and biochemical pathways that underlie variation in coloration in other amphibian groups including highly diverged aposematic taxa such as Australian myobatrachid frogs (e.g., *Pseudophryne corroboree)*, Malagasy poison frogs (Mantellidae), caecilians (e.g., *Schistometopum thomense*), and salamanders (e.g., *Salamandra salamandra*). Indeed, these methods have already enabled the identification of genes and loci involved in coloration in the salamander *S. salamandra bernardezi* [121].

Despite the numerous advances made with amphibian omics in elucidating evolutionary and ecological mechanisms, fully unraveling their genetic basis requires the generation of a vast number of genomes, given the comparative nature of these fields and the evolutionary uniqueness of each lineage. Some of the exciting research avenues in amphibians include behavioral adaptations like parental care [122, 123], milk production or skin feeding in caecilians [124, 125], spatial navigation [126]; adaptations to environmental conditions, like niche expansion due to the evolution of gliding ability [127], the evolution of lunglessness [128, 129] or predator-prey interactions like unusual defense mechanisms, such as the ability of some newts to pierce their ribs through toxin glands in their skin [130, 131].

### Conservation

Amphibians are the most endangered class of vertebrates with current estimates suggesting that more than 40% of species are threatened with extinction [132]. The threats amphibians face continue to increase [132], creating a clear need to develop innovative and effective methods to conserve them. Paradoxically, current rates of amphibian species description are exponential, and numerous candidate species are being flagged worldwide. This suggests that we are still far from overcoming the amphibian Linnean shortfall, especially in tropical regions [133, 134]. Hence, numbers of threatened species are likely underestimated, as undescribed species cannot be assessed and are more likely to become extinct [135]. Further, the conservation status of many amphibians remains unknown, especially for tropical species [136] and for a number of soil-dwelling caecilians for which only a limited number of specimens are available [137]. Generating genomic data is one method to address this challenge, as it can be used to estimate both evolutionary potential and extinction risk [138, 139]. Genomes are also vital for understanding species boundaries and the geographic distribution of genetic diversity within species, and for identifying populations under higher risk due to anthropogenic pressures or climate change [21, 22, 140, 141]. These features make genomic resources invaluable for developing species conservation action plans [142].

Amphibian conservation efforts should leverage population genetic theory and the burgeoning field of conservation genomics. These approaches enable the quantification of both neutral and adaptive diversity across genomes, thereby facilitating the promotion of adaptive potential or genetic rescue through translocation programs [143–146]. High quality genomes can also facilitate more comprehensive genomic diversity analyses, enabling the analyses of structural variants in addition to single nucleotide polymorphisms (SNPs), which are often overlooked, and an improve of the runs of homozygosity (ROH) analyses.

Typically, these studies begin with the genomic characterization of populations across various environmental conditions, assessing population genetic health and disease risk [147, 148]. They can also support monitoring and surveillance efforts by identifying populations most at risk of declines due to potential genetic threats like maladaptive alleles, genetic load, inbreeding and outbreeding depression, hybridization, and/or genetic incompatibility [143, 149]. Increased monitoring and maintenance of genomic diversity are key targets of many national and international recommendations such as the US Endangered Species Act [150], the Kunming-Montreal Global Biodiversity Monitoring Framework [151], and the Amphibian Conservation Action Plan[142].

A more specific application of amphibian genomics for conservation requires understanding the genetic basis of traits that impact fitness, such as disease resistance or climate change tolerance. The increased availability of long-read sequencing technology is particularly valuable in addressing the challenges of identifying highly variable gene regions accountable for immunological processes such as the major histocompatibility complex (MHC) [152]. This information can be used to promote adaptation using approaches like Targeted Genetic Intervention (TGI), which aims to increase the frequency of adaptive alleles with approaches such as selective breeding, genome editing, or targeted gene flow [153]. Considerable effort has been invested in understanding the genetic basis of resistance to the devastating amphibian disease chytridiomycosis. This has resulted in the identification of multiple candidate genes [154–156] that could be targeted to increase chytridiomycosis resistance with TGI.

Additionally, the efficacy of TGI at increasing chytridiomycosis resistance has already been demonstrated in North American mountain yellow-legged frogs (*Rana muscosa* and *R. sierrae*) where translocation of resistant individuals increased recipient population persistence [157]. Despite the obvious appeal of using genetic intervention approaches for conservation, these methods should be evaluated in contained facilities whenever possible and accompanied by long-term monitoring to ensure their efficacy and rule out any unintended impacts [153, 158–160]. Although such conservation interventions require extensive resources, this may be the only effective method for restoring some species to the wild, especially in those threatened by intractable threats such as chytridiomycosis [161].

## Challenges for amphibian genomic research and ways forward

The future of amphibian omics research will rely on high-quality reference genomes, which necessitates overcoming unique bioinformatic challenges in genome assembly and securing high-quality starting materials (e.g., tissue, blood). Additionally, challenges in obtaining funding, particularly in low-income countries, exacerbate these issues. Here, we outline these challenges in amphibian omics and highlight emerging developments aimed at addressing them.

The large genomes of amphibians increase requirements and costs for sequencing, computing, and data storage [6, 162]. Despite technological advancements and decreasing service costs, assembling these genomes remains methodologically challenging due to the notable intron lengths and repetitive content of amphibian genomes [163], especially when repeat lengths exceed sequencing read lengths. Regions of low complexity can result in erroneously joined contigs [164] or a significant loss of sequence information (by as much as 16%) through the collapsing of repetitive sequences [165]. Polyploidy has also evolved repeatedly in amphibians [166, 167], making haplotype-specific assemblies challenging and may require dramatically increased sequencing and computational efforts [168, 169]. The development of long-read sequencing (e.g. PacBio HiFi, ONT), optical mapping and 3C technology (i.e., Hi-C scaffolding) is therefore especially important for assembling amphibian genomes [164, 170].

Annotations are as crucial as genome assemblies, but current homology-based approaches using ortholog databases like UniProt [171] often miss or poorly annotate genes, especially polymorphic genes or those lacking representation in model taxa. This limits amphibian studies on gene evolution [72], repeats [16, 163], or immune genes [172].

Additionally, functional genomics tools like gene editing, *in vitro* fertilization and transgenesis are rare for most amphibians [153, 173], developed primarily in model species (e.g., *Xenopus* spp., *Ambystoma mexicanum*) [61, 174–177]. Immortal cell lines have been successfully generated for some amphibians [178] and protocols have been established to facilitate the initiation of spontaneously arising cell lines for a subset of anurans [179]. However, establishing cell cultures for most species requires extensive problem-solving and expertise [178].

Most tissue sampling protocols for sequencing reference genomes recommend harvesting samples from fresh tissue, followed immediately by flash freezing in liquid nitrogen (LN2) and storing at −80°C until extraction (https://www.vertebrategenomelab.org/resources/guidelines). This often requires fieldwork with many logistical challenges.

The small body sizes and blood volumes of most amphibians (e.g., < 30 g) may necessitate lethal sampling to obtain sufficient high-molecular-weight DNA for generating reference genomes (HMW, reaching 100 Kb or ultra HMW, reaching 1 Mb) [180, 181]. While this characteristic is shared with other taxonomic groups (e.g., invertebrates), lethal sampling may not always be legally permitted or ethically advisable in amphibians, especially for threatened species or those in captive collections [182]. Non-lethal sampling approaches, such as buccal swabs or toe or tail clips, are increasingly viable for various genomic applications, including low-coverage whole genome sequencing or targeted sequencing approaches [183, 184]. Until these become suitable for reference-grade genome sequencing, an alternative to minimize sampling impacts may be to use tadpoles instead of adults (e.g., to generate the genome of *Taudactylus pleione* [185]).

Working with museum or natural history collections [the burgeoning field of “museomics”; 186] is a promising avenue of research for circumventing the intrinsic problems of sample collection. Moreover, it allows access to past amphibian biodiversity and is revolutionizing amphibian taxonomy by integrating DNA from name-bearing type specimens, overcoming impediments like uncertainty in nomenclature, species complexes, and cryptic species [187–190]. Key challenges of such research include issues with DNA degradation, preservation methods, and contamination that need to be overcome [191–193]. This is particularly relevant for wet-preserved amphibian specimens, as retrieving DNA can be challenging due undocumented fixation and preservation methods that may alter nucleotide integrity. Methodological advances in laboratory protocols [e.g., 194, 195, 196] and the development of sequencing strategies, such as ‘Barcode Fishing’, have made significant progress in addressing these challenges, including the ability to sequence extinct species [187, 188, 197–199]. In the current era, even limited sequences from taxonomic type specimens are of unparalleled importance, especially for species identification using genetic data, by those applying methods like eDNA and metagenomics [200].

Other noteworthy challenges, that are not necessarily unique to amphibians, include securing collection and research permits, maintaining ultracold storage and an uninterrupted cold chain during transport, and adhering to regulations for the international movement of biological samples across political borders. [201, 202]. Amphibian-specific challenges, however, can arise due to their biological, ecological, and conservation characteristics. Centers of amphibian diversity and endemism include remote, highly specialized habitats, such as tropical montane forests, cave systems or isolated wetlands. Moreover, many amphibians have specialized aquatic, subterranean or arboreal ecologies, are mostly nocturnal and highly seasonal. These factors make fieldwork and sample transportation challenging, especially in regions with poor infrastructure, inadequate storage facilities, socio-political conflicts, and limited funding for research, conservation, and public awareness.

While eliminating some of these practical and political challenges in amphibian fieldwork is beyond the scope of individual researchers, the growing accessibility of genomic data calls for increased awareness of the principles of fair and equitable access to genetic resources, as outlined by the Convention on Biological Diversity (CBD) and further elaborated by the Nagoya Protocol https://www.cbd.int/abs/default.shtml). Indigenous peoples and local communities (IPLC) are often custodians of genetic resources (physical material) sought by researchers, requiring that all parties enter into collaborative and equitable agreements on access and benefit-sharing (ABS) before embarking on a genomics project [203–207].

## Aims, priorities, and structure of the Amphibian Genomics Consortium (AGC)

The AGC (https://mvs.unimelb.edu.au/amphibian-genomics-consortium) was launched in March 2023 to address the aforementioned knowledge gaps through technological advances and international cooperation. The mission of the AGC is to enhance international and interdisciplinary collaboration among amphibian researchers, expand amphibian genomic resources, and effectively utilize genomic data and functional resources to close the gap between genome biologists, scientists, and conservation practitioners. The leadership structure of the AGC consists of a director, two co-directors, and a 10-member board. The board was carefully chosen to ensure gender equality, diversity of scientific disciplines, career stages, and representation from various geographic regions.

The first actions of the AGC include hosting monthly regular meetings that showcase advances in amphibian genomics research, developing technical resources and best practices guidelines (through discussions facilitated in a Discord channel), improving amphibian genome annotation, supporting travel for students and early career researchers, hosting networking events at conferences, and conducting virtual and in-person computational workshops. Details of these activities can be found in the AGC website. The AGC plans to secure funding to sequence high-priority amphibian species (see The AGC’s genome sequencing targets section and Table 1).

Additionally, the AGC aims to facilitate amphibian sample collection for broader taxonomic consortia. The AGC is already affiliated with the Earth BioGenome Project (EBP; [208]) and AmphibiaWeb (https://amphibiaweb.org), reinforcing its commitment to advancing amphibian genomics and conservation efforts.

### AGC membership

At the time of the submission of this work, the AGC had 282 members from 41 countries (6 in Africa, 131 in the Americas, 27 in Asia, 29 in Australasia, and 89 in Europe), with membership continuing to increase (Fig. 4). Although the membership is geographically diverse, disparity persists across regions. The recruitment of members from underrepresented countries will be a key focus of the AGC, with a particular emphasis on regions known for high amphibian diversity and/or endemism such as Central and South America, and Southeast Asia. We promote equity between members by providing additional support and opportunities to those from developing countries and underrepresented groups. This includes eliminating membership fees, scheduling online meetings at alternating times to accommodate global time zones, facilitating discussion groups on the cloud-based collaboration platform Discord, and translating AGC correspondence into multiple languages. Furthermore, we are also committed to fostering knowledge and skills transfer to all emerging scientists worldwide, and we actively encourage early career researchers to join the initiative and participate in governance.

**Figure 4.**
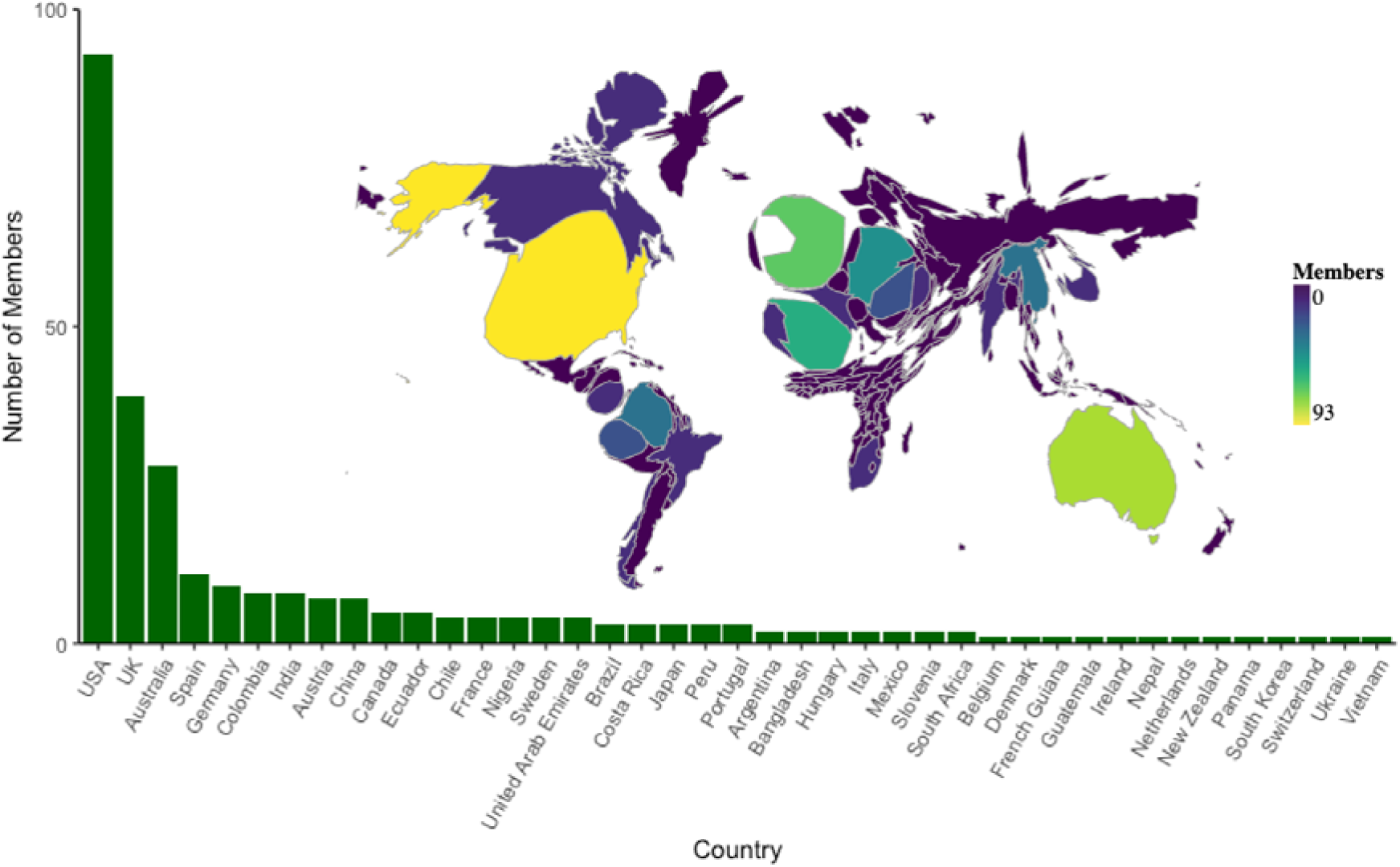
Amphibian Genomics Consortium (AGC) membership by country. Inset map showing the size of each country scaled by number of members in the AGC.

### Current use and perception of genomics technologies by members of the AGC

The AGC leadership designed a 23-question survey to investigate consortium members’ experiences in amphibian genomics (questions can be found in Supplementary Table S1). The survey was distributed using the Qualtrics XM platform and remained active from the 4^th^ of March to the 27^th^ of December 2023. We collected responses from a total of 133 AGC members from 32 countries with different expertise in sequencing approaches and bioinformatics techniques, who primarily work on the ecology and evolution of anurans. Overall, respondents emphasized the urgency of filling knowledge gaps in amphibian genomics due to the current conservation crisis, pinpointing the necessity to expand the number of high-quality chromosome-level amphibian genomes. Additionally, there was strong agreement among survey respondents that the generation of new genomic resources needs to be coupled with the improvement and accessibility of annotation processes. A better development of sharing computational expertise among members and resources internationally was also underscored. More than half of the survey participants said they use sequencing technologies for their studies (70 of the 133). About half of the respondents said their main work activities were “genomics lab work” or “computational analyses” (48% and 57%, respectively).

To evaluate consortium members’ experience in amphibian genomics, we applied a principal components analysis to the quantitative responses. Bioinformatic competencies and perceived challenges of the AGC respondents were grouped in two dimensions, respectively (Fig. 5A and Fig. S2; see Supplementary Material for methodological information). To explain the variation of these two new variables, we used the scientific expertise of AGC members, the funding success, and two variables related to the country of main affiliation of the respondent: the number of amphibian species and gross domestic expenditure on R&D (GERD) per capita, as explanatory variables. Amphibian genomics expertise and identified challenges varied substantially among respondents. The number of amphibian species and GERD per capita of the respondent’s main affiliation country did not capture this variation (Fig. 5B and Fig. S3; see Supplementary Material for methodological information). Instead, genomics funding success and years of scientific expertise were, as expected, positively correlated and both variables were associated with a reduction in the perceived challenges associated with amphibian genomics.

**Figure 5.**
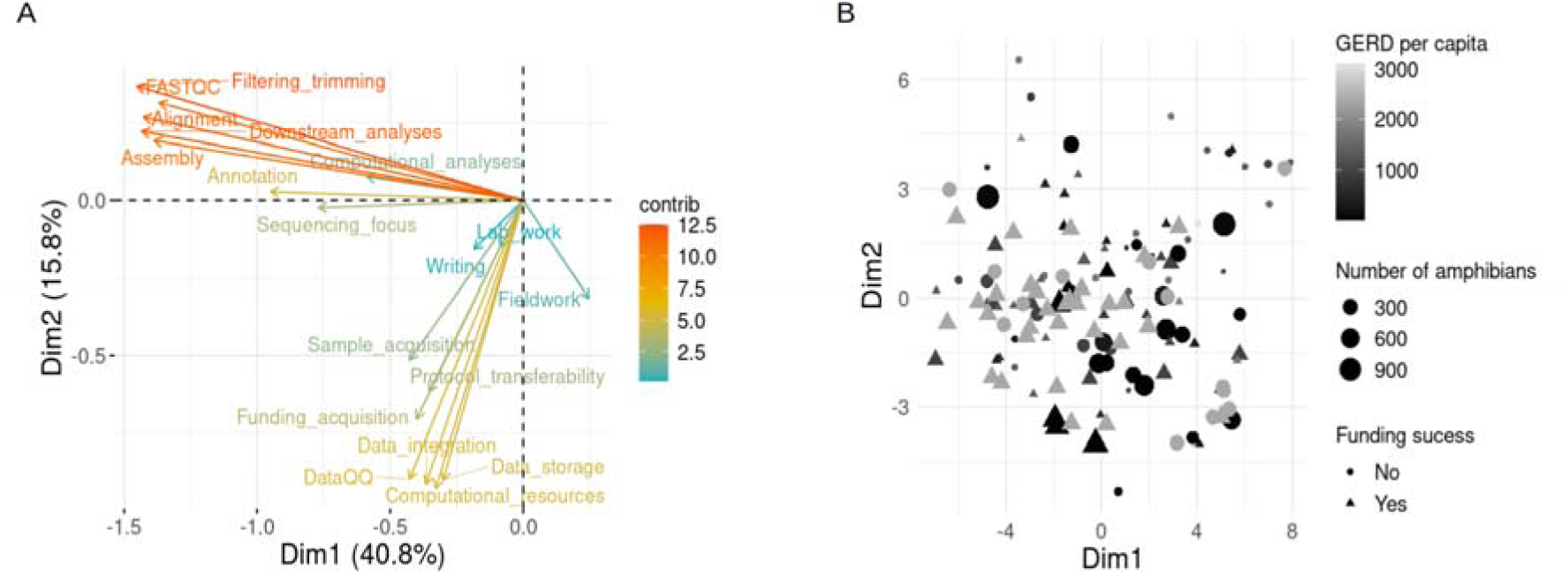
Sequencing competencies and identified challenges of the members of the Amphibian Genomics Consortium (AGC). (A) Representation of the contribution of the AGC survey quantitative questions to the first dimensions after computing a principal component analysis (PCA). Bioinformatic competencies and perceived challenges were grouped into dimensions one and two, respectively. (B) Scatter plot showing PCA scores for each AGC survey respondent. Respondent answers are coded by the qualitative question about funding success for amphibian genomics projects using shape; number of amphibian species of the respondent main affiliation country by size, and gross domestic expenditure on R&D (GERD) per capita of the respondent main affiliation country by gray-scale color coded. Information about the number of amphibian species per country was obtained from AmphibiaWeb. GERD per capita was calculated using information from the UNESCO and World Bank websites from the information about the most recent year for each country.

### The AGC’s genome sequencing targets

Following the efforts of genomics consortia for other tetrapod groups [e.g., 209], and previous research [22], we identified 48 amphibian families for which no representative genomes had been sequenced and selected one representative species from each family for our sequencing priority list (Fig. 2B and Table 1). We propose 48 candidate species based on their IUCN Red List category, ecological and evolutionary distinctiveness, and the availability of other genomics records, especially transcriptomics. This list includes 38 anurans, four urodeles, and six caecilians.

We recommend this priority list as a starting point. If suitable sample material from other species within the targeted families becomes available, those species could replace the ones currently proposed. fAdditionally, we aim to build upon the efforts of existing genomics consortia such as the Vertebrate Genomes Project (VGP), hence, we included two species with draft genomes in the GenomeArk (https://www.genomeark.org/) in our sequencing target list.

### The AGC’s stance on resource and benefits sharing

With increasingly easy access to genomic data, researchers and industry need to be aware of the principles of fair and equitable access to genetic resources, as stipulated by Convention on Biological Diversity (CBD) and expanded upon by the Nagoya Protocol (https://www.cbd.int/abs/default.shtml). As a negative example from amphibians, *Phyllomedusa bicolor* skin secretions traditionally used by Amazonian Indigenous peoples were patented by actors in the US, Japan, Russia and elsewhere, promoting the ‘legal’ but unfair appropriation of genetic resources and potentially the traditional knowledge itself from the Matses and other Indigenous tribes [210].

To promote better practices, researchers should allocate the necessary time and funds for prior consultation during fieldwork planning and seek guidance from their National Focal Points on ABS. How the concept of ABS may be applied to the downstream use of the digital sequence information (DSI) generated has yet to be resolved. However, there are currently developments underway that may provide a solution (https://www.cbd.int/dsi-gr). It is imperative that this issue be considered going forward [see for example 211]. Moreover, voucher specimens and duplicate tissue samples should be deposited in local natural history collections or preferred partners of the local communities [212, 213].

The global genomics community should strive to ensure that sequencing projects occur within the country of origin of the samples and discourage ‘parachute’ or ‘helicopter science’ [214, 215]. Oxford Nanopore Technology (ONT) may be promising solution, providing comparatively affordable access to equipment and reagents for ultra-long read sequencing that can even be done directly in the field [216]. However, optimization for non-model organisms, along with the startup costs for this infrastructure remain prohibitive for many scientists from low-income countries. Moving forward, the goal should be to apply these technologies in collaboration with local researchers. Programs like the In Situ Laboratories Initiative (https://insitulabs.org/hubs/) aim to overcome these challenges by providing affordable access to high-tech laboratories in remote biodiverse areas. Such collaborative projects should proceed from finding shared interests, developing ideas, realizing the shared benefits from research outputs, and focusing on capacity-building efforts [217].

## Conclusion and call to action

Moving forward, the AGC is committed to supporting amphibian sequencing initiatives worldwide, with a particular emphasis on taxonomic groups lacking representation, and species from biodiverse countries within a conservation framework (Table 1). Local sequencing initiatives will be given priority whenever feasible to promote the development of *in situ* research efforts and facilities. We will achieve this goal building strong networks between researchers and conservation practitioners and by providing an open list of members and their expertise. Additionally, we aim to provide funding and training opportunities to facilitate collaboration among underrepresented groups, molecular and organismal biologists, bioinformaticians, and conservation practitioners. We will also support the development of a concept of Access and Benefit Sharing policies that can be applied to the downstream use of the digital sequence information (DSI), including long-term storage and access. Further, the AGC aspires to stimulate public and scientific interest in amphibian research and, ultimately, to enhance conservation outcomes for this intriguing and highly endangered group of vertebrates.

We hope that the recent advancements in technology, a focus on equitable research, and the integration of the research community to form the AGC will ignite research to revolutionize amphibian conservation and our understanding of their fascinating biology, ecology and evolution. By addressing the challenges outlined, supporting and promoting amphibian genomics research and uniting amphibian researchers worldwide, the AGC aims to fill the huge gap in genomic data for this diverse group of tetrapods and in doing so, propel amphibian genomics research into the future.

## List of Abbreviations

ABS: access and benefit-sharing
AGC: Amphibian Genomics Consortium
CBD: Convention on Biological Diversity
DSI: digital sequence information
EBP: Earth BioGenome Project
GERD: gross domestic expenditure on research and development
GoaT: Genomes on a Tree
HMW: High molecular weight DNA
IPLC: Indigenous peoples and local communities
IUCN: International Union for Conservation of Nature
ONT: Oxford Nanopore Technology
VGP: Vertebrate Genomes Project

## Declarations

### Ethics approval and consent to participate

Not applicable.

### Consent for publication

Not applicable.

### Availability of data and materials

Not applicable.

### Authors’ contributions

T.A.K. and M.T.-S. drafted the manuscript. T.A.K., M.T.-S., H.C.L., K.S., M.H.Y., S.T.M., A.J.C. contributed text to the first draft, M.T.-S. and T.A.K. analyzed the data and created the figures, members of the Amphibian Genomics Consortium (AGC) reviewed later drafts. T.A.K., M.T.-S, C.D., N.J.F, Y.C., R.D.T., H.C.L, V.L.N.A., R.M., K.S.-S., H.M., K.C.W.V, M.H.Y., and M.S. contributed to revising the manuscript for resubmission. All authors reviewed the manuscript.

## Acknowledgments

Contributing members of the Amphibian Genomics Consortium (AGC) in alphabetical order: Aldemar A. Acevedo, Steven J. R. Allain, Lisa N. Barrow, M. Delia Basanta, Roberto Biello, Gabriela B. Bittencourt-Silva, Amaël Borzée, Ian G. Brennan, Rafe M. Brown, Natalie Calatayud, Hugo Cayuela, Jing Chai, Ignacio De la Riva, Lana J. Deaton, Khalid A. E. Eisawi, Kathryn R. Elmer, W. Chris Funk, Giussepe Gagliardi-Urrutia, Wei Gao, Mark J. Goodman, Sandra Goutte, Melissa Hernandez Poveda, Tomas Hrbek, Oluyinka A. Iyiola, Gregory F.M. Jongsma, J. Scott Keogh, Tianming Lan, Pablo Lechuga-Paredes, Emily Moriarty Lemmon, Stephen C. Lougheed, Thom A. Lyons, Mariana L. Lyra, Jimmy A. McGuire, Marco A. Mendez, Hosne Mobarak, Edina Nemesházi, Tao T. Nguyen, Michaël P.J. Nicolaï, Lotanna M. Nneji, John B. Owens, Hibraim Pérez-Mendoza, Nicolas Pollet, Megan L. Power, Mizanur Rahman, Hans Recknagel, Ariel Rodríguez, Santiago R. Ron, Joana Sabino-Pinto, Yongming Sang, Suman Sapkota, Rosio G. Schneider, Laura Schulte, Ana Serra Silva, Lee F. Skerratt, Nicholas Strowbridge, Karthikeyan Vasudevan, Govindappa Venu, Lucas Vicuña, David R. Vieites, Judit Vörös, Matt West, Mark Wilkinson, Guinevere O. U. Wogan. We also appreciate the helpful suggestions from Jia-Tang Li, Jun Li, Wei Wu, and Hua Wu.

## References

1. Hellsten U, Harland RM, Gilchrist MJ, Hendrix D, Jurka J, Kapitonov V, Ovcharenko I, Putnam NH, Shu S, Taher L et al: The Genome of the Western Clawed Frog*Xenopus tropicalis*. Science 2010, 328(5978):633–636.

2. Band MR, Larson JH, Rebeiz M, Green CA, Heyen DW, Donovan J, Windish R, Steining C, Mahyuddin P, Womack JE et al: An ordered comparative map of the cattle and human genomes. Genome Res 2000, 10(9):1359–1368.

3. Aparicio S, Chapman J, Stupka E, Putnam N, Chia JM, Dehal P, Christoffels A, Rash S, Hoon S, Smit A et al: Whole-genome shotgun assembly and analysis of the genome of *Fugu rubripes*. Science 2002, 297(5585):1301–1310.

4. Collins FS, Green ED, Guttmacher AE, Guyer MS, on behalf of the USNHGRI: A vision for the future of genomics research. Nature 2003, 422(6934):835–847.

5. Hillier LW, Miller W, Birney E, Warren W, Hardison RC, Ponting CP, Bork P, Burt DW, Groenen MAM, Delany ME et al: Sequence and comparative analysis of the chicken genome provide unique perspectives on vertebrate evolution. Nature 2004, 432(7018):695–716.

6. Stöck M, Kratochvíl L, Kuhl H, Rovatsos M, Evans BJ, Suh A, Valenzuela N, Veyrunes F, Zhou Q, Gamble T et al: A brief review of vertebrate sex evolution with a pledge for integrative research: towards ‘sexomics’. Philosophical Transactions of the Royal Society B: Biological Sciences 2021, 376(1832):20200426.

7. Pillay R, Venter M, Aragon-Osejo J, González-del-Pliego P, Hansen AJ, Watson JE, Venter O: Tropical forests are home to over half of the world’s vertebrate species. Front Ecol Environ 2022, 20(1):10–15.

8. Gregory TR: The evolution of the genome: Elsevier; 2011.

9. Biscotti MA, Carducci F, Olmo E, Canapa A: Vertebrate Genome Size and the Impact of Transposable Elements in Genome Evolution. In: Evolution, Origin of Life, Concepts and Methods. Edited by Pontarotti P. Cham: Springer International Publishing; 2019: 233-251.

10. Weisrock DW, Hime PM, Nunziata SO, Jones KS, Murphy MO, Hotaling S, Kratovil JD: Surmounting the large-genome “problem” for genomic data generation in salamanders. Population genomics: wildlife 2021:115–142.

11. Schartl M, Woltering JM, Irisarri I, Du K, Kneitz S, Pippel M, Brown T, Franchini P, Li J, Li M et al: The genomes of all lungfish inform on genome expansion and tetrapod evolution. Nature 2024.

12. Liedtke HC, Gower DJ, Wilkinson M, Gomez-Mestre I: Macroevolutionary shift in the size of amphibian genomes and the role of life history and climate. Nature Ecology & Evolution 2018, 2(11):1792–1799.

13. Nowoshilow S, Schloissnig S, Fei J-F, Dahl A, Pang AWC, Pippel M, Winkler S, Hastie AR, Young G, Roscito JG et al: The axolotl genome and the evolution of key tissue formation regulators. Nature 2018.

14. Brown T, Elewa A, Iarovenko S, Subramanian E, Araus AJ, Petzold A, Susuki M, Suzuki K-iT, Hayashi T, Toyoda A et al: Sequencing and chromosome-scale assembly of the giant *Pleurodeles waltl* genome. bioRxiv 2022:2022.2010.2019.512763.

15. Talavera A, Palmada-Flores M, Burriel-Carranza B, Valbuena-Ureña E, Mochales-Riaño G, Adams DC, Tejero-Cicuéndez H, Soler-Membrives A, Amat F, Guinart D et al: Genomic insights into the Montseny brook newt (*Calotriton arnoldi*), a Critically Endangered glacial relict. iScience 2024, 27(1):108665.

16. Kosch TA, Crawford AJ, Mueller RL, Wollenberg Valero KC, Power ML, Rodríguez A, O’Connell LA, Young ND, Skerratt LF: Comparative analysis of amphibian genomes: an emerging resource for basic and applied research. bioRxiv 2024:2023.2002.2027.530355.

17. Sun C, Shepard DB, Chong RA, López Arriaza J, Hall K, Castoe TA, Feschotte C, Pollock DD, Mueller RL: LTR retrotransposons contribute to genomic gigantism in plethodontid salamanders. Genome Biol Evol 2012, 4(2):168–183.

18. Fu T-T, Sun Y-B, Gao W, Long C-B, Yang C-H, Yang X-W, Zhang Y, Lan X-Q, Huang S, Jin J-Q et al: The highest-elevation frog provides insights into mechanisms and evolution of defenses against high UV radiation. Proceedings of the National Academy of Sciences 2022, 119(46):e2212406119.

19. Challis R, Kumar S, Sotero-Caio C, Brown M, Blaxter M: Genomes on a Tree (GoaT): A versatile, scalable search engine for genomic and sequencing project metadata across the eukaryotic tree of life [version 1; peer review: 2 approved]. Wellcome Open Research 2023, 8(24).

20. Gregory TR: Animal genome size database. 2024.

21. Calboli FCF, Fisher MC, Garner TWJ, Jehle R: The need for jumpstarting amphibian genome projects. Trends Ecol Evol 2011, 26(8):378–379.

22. Funk WC, Zamudio KR, Crawford AJ: Advancing Understanding of Amphibian Evolution, Ecology, Behavior, and Conservation with Massively Parallel Sequencing. In: Population Genomics: Wildlife. Edited by Hohenlohe PA, Rajora OP. Cham: Springer International Publishing; 2018: 211–254.

23. Sun Y-B, Zhang Y, Wang K: Perspectives on studying molecular adaptations of amphibians in the genomic era. Zool Res 2020, 41(4):351.

24. Callery EM: There’s more than one frog in the pond: A survey of the Amphibia and their contributions to developmental biology. In: Semin Cell Dev Biol: 2006: Elsevier; 2006: 80–92.

25. Weaver C, Kimelman D: Move it or lose it: axis specification in *Xenopus*. Development 2004, 131(15):3491–3499.

26. Burggren WW, Warburton S: Amphibians as animal models for laboratory research in physiology. ILAR journal 2007, 48(3):260–269.

27. Naert T, Van Nieuwenhuysen T, Vleminckx K: TALENs and CRISPR/Cas9 fuel genetically engineered clinically relevant *Xenopus tropicalis* tumor models. Genesis 2017, 55(1-2):e23005.

28. Guille M, Grainger R: Genetics and Gene Editing Methods in *Xenopus laevis* and *Xenopus tropicalis*. Cold Spring Harbor Protocols 2023, 2023(6):pdb. top107045.

29. Horb M, Wlizla M, Abu-Daya A, McNamara S, Gajdasik D, Igawa T, Suzuki A, Ogino H, Noble A, France CdRBXti: *Xenopus* resources: transgenic, inbred and mutant animals, training opportunities, and web-based support. Frontiers in physiology 2019, 10:387.

30. Fisher M, James-Zorn C, Ponferrada V, Bell AJ, Sundararaj N, Segerdell E, Chaturvedi P, Bayyari N, Chu S, Pells T: Xenbase: key features and resources of the*Xenopus* model organism knowledgebase. Genetics 2023, 224(1):iyad018.

31. Liedtke HC, Wiens JJ, Gomez-Mestre I: The evolution of reproductive modes and life cycles in amphibians. Nat Commun 2022, 13(1):7039.

32. Bredeson JV, Mudd AB, Medina-Ruiz S, Mitros T, Smith OK, Miller KE, Lyons JB, Batra SS, Park J, Berkoff KC et al: Conserved chromatin and repetitive patterns reveal slow genome evolution in frogs. Nat Commun 2024, 15(1):579.

33. Liedtke HC, Harney E, Gomez-Mestre I: Cross-species transcriptomics uncovers genes underlying genetic accommodation of developmental plasticity in spadefoot toads. Mol Ecol 2021, 30(10):2220–2234.

34. Isdaner AJ, Levis NA, Pfennig DW: Comparative transcriptomics reveals that a novel form of phenotypic plasticity evolved via lineage-specific changes in gene expression. Ecol Evol 2023, 13(10):e10646.

35. Nemesházi E, Bókony V: HerpSexDet: the herpetological database of sex determination and sex reversal. Scientific Data 2023, 10(1):377.

36. Ma W-J, Veltsos P: The Diversity and Evolution of Sex Chromosomes in Frogs. Genes 2021, 12(4):483.

37. Schartl M, Schmid M, Nanda I: Dynamics of vertebrate sex chromosome evolution: from equal size to giants and dwarfs. Chromosoma 2016, 125:553–571.

38. Roco ÁS, Olmstead AW, Degitz SJ, Amano T, Zimmerman LB, Bullejos M: Coexistence of Y, W, and Z sex chromosomes in*Xenopus tropicalis*. Proceedings of the National Academy of Sciences 2015, 112(34):E4752–E4761.

39. Jeffries DL, Lavanchy G, Sermier R, Sredl MJ, Miura I, Borzée A, Barrow LN, Canestrelli D, Crochet P-A, Dufresnes C et al: A rapid rate of sex-chromosome turnover and non-random transitions in true frogs. Nat Commun 2018, 9(1):4088.

40. Cauret CM, Jordan DC, Kukoly LM, Burton SR, Anele EU, Kwiecien JM, Gansauge M-T, Senthillmohan S, Greenbaum E, Meyer M: Functional dissection and assembly of a small, newly evolved, W chromosome-specific genomic region of the African clawed frog *Xenopus laevis*. PLoS Genet 2023, 19(10):e1010990.

41. Kuhl H, Tan WH, Klopp C, Kleiner W, Koyun B, Ciorpac M, Feron R, Knytl M, Kloas W, Schartl M et al: A candidate sex determination locus in amphibians which evolved by structural variation between X- and Y-chromosomes. Nat Commun 2024, 15(1):4781.

42. Bertola LV, Hoskin CJ, Jones DB, Zenger KR, McKnight DT, Higgie M: The first linkage map for Australo-Papuan Treefrogs (family: Pelodryadidae) reveals the sex-determination system of the Green-eyed Treefrog (*Litoria serrata*). Heredity 2023, 131(4):263–272.

43. Bogart JP, Bi K, Fu J, Noble DW, Niedzwiecki J: Unisexual salamanders (genus *Ambystoma*) present a new reproductive mode for eukaryotes. Genome 2007, 50(2):119–136.

44. McElroy KE, Denton RD, Sharbrough J, Bankers L, Neiman M, Gibbs HL: Genome expression balance in a triploid trihybrid vertebrate. Genome Biol Evol 2017, 9(4):968–980.

45. Li Y, Ren Y, Zhang D, Jiang H, Wang Z, Li X, Rao D: Chromosome-level assembly of the mustache toad genome using third-generation DNA sequencing and Hi-C analysis. GigaScience 2019, 8(9):giz114.

46. Mikó Z, Nemesházi E, Ujhegyi N, Verebélyi V, Ujszegi J, Kásler A, Bertalan R, Vili N, Gál Z, Hoffmann OI: Sex reversal and ontogeny under climate change and chemical pollution: are there interactions between the effects of elevated temperature and a xenoestrogen on early development in agile frogs? Environ Pollut 2021, 285:117464.

47. Das B, Cai L, Carter MG, Piao Y-L, Sharov AA, Ko MS, Brown DD: Gene expression changes at metamorphosis induced by thyroid hormone in*Xenopus laevis* tadpoles. Dev Biol 2006, 291(2):342–355.

48. Schott RK, Bell RC, Loew ER, Thomas KN, Gower DJ, Streicher JW, Fujita MK: Transcriptomic evidence for visual adaptation during the aquatic to terrestrial metamorphosis in leopard frogs. BMC Biol 2022, 20(1):138.

49. Wollenberg Valero KC, Garcia-Porta J, Rodríguez A, Arias M, Shah A, Randrianiaina RD, Brown JL, Glaw F, Amat F, Künzel S et al: Transcriptomic and macroevolutionary evidence for phenotypic uncoupling between frog life history phases. Nat Commun 2017, 8(1):15213.

50. Zhao L, Liu L, Wang S, Wang H, Jiang J: Transcriptome profiles of metamorphosis in the ornamented pygmy frog *Microhyla fissipes* clarify the functions of thyroid hormone receptors in metamorphosis. Sci Rep 2016, 6(1):27310.

51. Palacios-Martinez J, Caballero-Perez J, Espinal-Centeno A, Marquez-Chavoya G, Lomeli H, Salas-Vidal E, Schnabel D, Chimal-Monroy J, Cruz-Ramirez A: Multi-organ transcriptomic landscape of *Ambystoma velasci* metamorphosis. Dev Biol 2020, 466(1-2):22–35.

52. Sanchez E, Küpfer E, Goedbloed DJ, Nolte AW, Lüddecke T, Schulz S, Vences M, Steinfartz S: Morphological and transcriptomic analyses reveal three discrete primary stages of postembryonic development in the common fire salamander, *Salamandra salamandra* . Journal of Experimental Zoology Part B: Molecular and Developmental Evolution 2018, 330(2):96–108.

53. Goedert D, Calsbeek R: Experimental evidence that metamorphosis alleviates genomic conflict. The American Naturalist 2019, 194(3):356–366.

54. Kyono Y, Raj S, Sifuentes CJ, Buisine N, Sachs L, Denver RJ: DNA methylation dynamics underlie metamorphic gene regulation programs in*Xenopus* tadpole brain. Dev Biol 2020, 462(2):180–196.

55. Pfennig KS: Facultative mate choice drives adaptive hybridization. Science 2007, 318(5852):965–967.

56. Levis NA, Pfennig DW: Innovation and diversification via plasticity-led evolution. In: Phenotypic plasticity & evolution. CRC Press; 2021: 211–240.

57. Pfennig DW, Collins JP: Kinship affects morphogenesis in cannibalistic salamanders. Nature 1993, 362(6423):836–838.

58. Pfennig DW, Reeve HK, Sherman PW: Kin recognition and cannibalism in spadefoot toad tadpoles. Anim Behav 1993, 46(1):87–94.

59. DeVore JL, Crossland MR, Shine R, Ducatez S: The evolution of targeted cannibalism and cannibal-induced defenses in invasive populations of cane toads. Proceedings of the National Academy of Sciences 2021, 118(35):e2100765118.

60. Brockes JP, Kumar A: Comparative aspects of animal regeneration. Annu Rev Cell Dev Biol 2008, 24:525–549.

61. Cox BD, Yun MH, Poss KD: Can laboratory model systems instruct human limb regeneration? Development 2019, 146(20):dev181016.

62. Torres-Sánchez M: Variation under domestication in animal models: the case of the Mexican axolotl. BMC Genomics 2020, 21(1):827.

63. Yu Q, Gates PB, Rogers S, Mikicic I, Elewa A, Salomon F, Lachnit M, Caldarelli A, Flores-Rodriguez N, Cesare AJ et al: Telomerase-independent maintenance of telomere length in a vertebrate. bioRxiv 2022:2022.2003.2025.485759.

64. Bruckskotten M, Looso M, Reinhardt R, Braun T, Borchardt T: Newt-omics: a comprehensive repository for omics data from the newt*Notophthalmus viridescens*. Nucleic Acids Res 2012, 40(Database issue):D895–900.

65. Feng Y-J, Blackburn DC, Liang D, Hillis DM, Wake DB, Cannatella DC, Zhang P: Phylogenomics reveals rapid, simultaneous diversification of three major clades of Gondwanan frogs at the Cretaceous–Paleogene boundary. Proceedings of the National Academy of Sciences 2017, 114(29):E5864–E5870.

66. Schott RK, Fujita MK, Streicher JW, Gower DJ, Thomas KN, Loew ER, Bamba Kaya AG, Bittencourt-Silva GB, Guillherme Becker C, Cisneros-Heredia D: Diversity and Evolution of Frog Visual Opsins: Spectral Tuning and Adaptation to Distinct Light Environments. Mol Biol Evol 2024, 41(4):msae049.

67. Rancilhac L, Irisarri I, Angelini C, Arntzen JW, Babik W, Bossuyt F, Künzel S, Lüddecke T, Pasmans F, Sanchez E et al: Phylotranscriptomic evidence for pervasive ancient hybridization among Old World salamanders. Mol Phylogen Evol 2021, 155:106967.

68. Dubey S, Maddalena T, Bonny L, Jeffries DL, Dufresnes C: Population genomics of an exceptional hybridogenetic system of*Pelophylax* water frogs. BMC Evol Biol 2019, 19(1):164.

69. Dufresnes C, Ambu J, Galán P, Sequeira F, Viesca L, Choda M, Álvarez D, Alard B, Suchan T, Künzel S: Delimiting phylogeographic diversity in the genomic era: application to an Iberian endemic frog. Zool J Linn Soc 2023:zlad170.

70. Ovchinnikov V, Uliano-Silva M, Wilkinson M, Wood J, Smith M, Oliver K, Sims Y, Torrance J, Suh A, McCarthy SA et al: Caecilian Genomes Reveal the Molecular Basis of Adaptation and Convergent Evolution of Limblessness in Snakes and Caecilians. Mol Biol Evol 2023, 40(5).

71. Jetz W, Pyron RA: The interplay of past diversification and evolutionary isolation with present imperilment across the amphibian tree of life. Nature Ecology & Evolution 2018, 2(5):850–858.

72. Siu-Ting K, Torres-Sánchez M, San Mauro D, Wilcockson D, Wilkinson M, Pisani D, O’Connell MJ, Creevey CJ: Inadvertent Paralog Inclusion Drives Artifactual Topologies and Timetree Estimates in Phylogenomics. Mol Biol Evol 2019, 36(6):1344–1356.

73. Hime PM, Lemmon AR, Lemmon ECM, Prendini E, Brown JM, Thomson RC, Kratovil JD, Noonan BP, Pyron RA, Peloso PL: Phylogenomics reveals ancient gene tree discordance in the amphibian tree of life. Syst Biol 2021, 70(1):49–66.

74. Portik DM, Streicher JW, Blackburn DC, Moen DS, Hutter CR, Wiens JJ: Redefining Possible: Combining Phylogenomic and Supersparse Data in Frogs. Mol Biol Evol 2023, 40(5).

75. Sun Y-B, Xiong Z-J, Xiang X-Y, Liu S-P, Zhou W-W, Tu X-L, Zhong L, Wang L, Wu D-D, Zhang B-L et al: Whole-genome sequence of the Tibetan frog *Nanorana parkeri* and the comparative evolution of tetrapod genomes. Proceedings of the National Academy of Sciences 2015, 112(11):E1257–E1262.

76. Torres-Sánchez M, Gower DJ, Alvarez-Ponce D, Creevey CJ, Wilkinson M, San Mauro D: What lies beneath? Molecular evolution during the radiation of caecilian amphibians. BMC Genomics 2019, 20(1):354.

77. Wellenreuther M, Mérot C, Berdan E, Bernatchez L: Going beyond SNPs: The role of structural genomic variants in adaptive evolution and species diversification. Mol Ecol 2019, 28(6):1203–1209.

78. Li J-T, Gao Y-D, Xie L, Deng C, Shi P, Guan M-L, Huang S, Ren J-L, Wu D-D, Ding L: Comparative genomic investigation of high-elevation adaptation in ectothermic snakes. Proceedings of the National Academy of Sciences 2018, 115(33):8406–8411.

79. Storz JF: High-Altitude Adaptation: Mechanistic Insights from Integrated Genomics and Physiology. Mol Biol Evol 2021, 38(7):2677–2691.

80. Yu L, Wang G-D, Ruan J, Chen Y-B, Yang C-P, Cao X, Wu H, Liu Y-H, Du Z-L, Wang X-P: Genomic analysis of snub-nosed monkeys (*Rhinopithecus*) identifies genes and processes related to high-altitude adaptation. Nat Genet 2016, 48(8):947–952.

81. Seimon TA, Seimon A, Daszak P, Halloy SR, Schloegel LM, Aguilar CA, Sowell P, Hyatt AD, Konecky B, E Simmons J: Upward range extension of Andean anurans and chytridiomycosis to extreme elevations in response to tropical deglaciation. Global Change Biol 2007, 13(1):288–299.

82. Acosta-Galvis AR: Ranas, salamandras y caecilias (Tetrapoda: Amphibia) de Colombia. Biota colombiana 2000, 1(3).

83. Yang W, Qi Y, Fu J: Genetic signals of high-altitude adaptation in amphibians: a comparative transcriptome analysis. BMC Genet 2016, 17(1):134.

84. Lu B, Jin H, Fu J: Molecular convergent and parallel evolution among four high-elevation anuran species from the Tibetan region. BMC Genomics 2020, 21:1–14.

85. Cayuela H, Dorant Y, Forester BR, Jeffries DL, Mccaffery RM, Eby LA, Hossack BR, Gippet JM, Pilliod DS, Chris Funk W: Genomic signatures of thermal adaptation are associated with clinal shifts of life history in a broadly distributed frog. J Anim Ecol 2022, 91(6):1222–1238.

86. Wang G-D, Zhang B-L, Zhou W-W, Li Y-X, Jin J-Q, Shao Y, Yang H-c, Liu Y-H, Yan F, Chen H-M: Selection and environmental adaptation along a path to speciation in the Tibetan frog *Nanorana parkeri*. Proc Natl Acad Sci USA 2018, 115(22):E5056–E5065.

87. Sun Y-B, Fu T-T, Jin J-Q, Murphy RW, Hillis DM, Zhang Y-P, Che J: Species groups distributed across elevational gradients reveal convergent and continuous genetic adaptation to high elevations. Proceedings of the National Academy of Sciences 2018, 115(45):E10634–E10641.

88. Hutchison VH, Haines HB, Engbretson G: Aquatic life at high altitude: Respiratory adaptations in the lake titicaca frog, *Telmatobius culeus*. Respiration Physiology 1976, 27(1):115–129.

89. Dunn ER: The salamanders of the family Plethodontidae, vol. 7: Smith College; 1926.

90. Daly J, Garraffo H, Pannell L, Spande T, Severini C, Erspamer V: Alkaloids from Australian frogs (Myobatrachidae): pseudophrynamines and pumiliotoxins. J Nat Prod 1990, 53(2):407–421.

91. Darst CR, Cummings ME: Predator learning favours mimicry of a less-toxic model in poison frogs. Nature 2006, 440(7081):208–211.

92. Hayes RA, Piggott AM, Dalle K, Capon RJ: Microbial biotransformation as a source of chemical diversity in cane toad steroid toxins. Bioorganic & Medicinal Chemistry Letters 2009, 19(6):1790–1792.

93. Tóth Z, Kurali A, Móricz ÁM, Hettyey A: Changes in Toxin Quantities Following Experimental Manipulation of Toxin Reserves in*Bufo bufo* Tadpoles. J Chem Ecol 2019, 45(3):253–263.

94. Vaelli PM, Theis KR, Williams JE, O’Connell LA, Foster JA, Eisthen HL: The skin microbiome facilitates adaptive tetrodotoxin production in poisonous newts. eLife 2020, 9:e53898.

95. Daly JW, Martin Garraffo H, Spande TF, Jaramillo C, Stanley Rand A: Dietary source for skin alkaloids of poison frogs (Dendrobatidae)? J Chem Ecol 1994, 20:943–955.

96. Darst CR, Menéndez-Guerrero PA, Coloma LA, Cannatella DC: Evolution of dietary specialization and chemical defense in poison frogs (Dendrobatidae): a comparative analysis. The American Naturalist 2005, 165(1):56–69.

97. Caty SN, Alvarez-Buylla A, Byrd GD, Vidoudez C, Roland AB, Tapia EE, Budnik B, Trauger SA, Coloma LA, O’Connell LA: Molecular physiology of chemical defenses in a poison frog. J Exp Biol 2019, 222(12):jeb204149.

98. Alvarez-Buylla A, Fischer M-T, Moya Garzon MD, Rangel AE, Tapia EE, Tanzo JT, Soh HT, Coloma LA, Long JZ, O’Connell LA: Binding and sequestration of poison frog alkaloids by a plasma globulin. eLife 2023, 12:e85096.

99. Abderemane-Ali F, Rossen ND, Kobiela ME, Craig RA, Garrison CE, Chen Z, Colleran CM, O’Connell LA, Du Bois J, Dumbacher JP: Evidence that toxin resistance in poison birds and frogs is not rooted in sodium channel mutations and may rely on “toxin sponge” proteins. J Gen Physiol 2021, 153(9):e202112872.

100. Márquez R, Ramírez-Castañeda V, Amézquita A: Does batrachotoxin autoresistance coevolve with toxicity in *Phyllobates* poison-dart frogs? Evolution 2019, 73(2):390–400.

101. Tarvin RD, Borghese CM, Sachs W, Santos JC, Lu Y, O’Connell LA, Cannatella DC, Harris RA, Zakon HH: Interacting amino acid replacements allow poison frogs to evolve epibatidine resistance. Science 2017, 357(6357):1261–1266.

102. Tarvin RD, Santos JC, O’Connell LA, Zakon HH, Cannatella DC: Convergent substitutions in a sodium channel suggest multiple origins of toxin resistance in poison frogs. Mol Biol Evol 2016, 33(4):1068–1080.

103. Shibao PYT, Cologna CT, Morandi-Filho R, Wiezel GA, Fujimura PT, Ueira-Vieira C, Arantes EC: Deep sequencing analysis of toad *Rhinella schneideri* skin glands and partial biochemical characterization of its cutaneous secretion. J Venom Anim Tox incl Trop Dis 2018, 24(1):36.

104. Torres-Sánchez M, Wilkinson M, Gower DJ, Creevey CJ, San Mauro D: Insights into the skin of caecilian amphibians from gene expression profiles. BMC Genomics 2020, 21(1):515.

105. Liscano Martinez Y, Arenas Gómez CM, Smith J, Delgado JP: A tree frog (*Boana pugnax*) dataset of skin transcriptome for the identification of biomolecules with potential antimicrobial activities. Data in Brief 2020, 32:106084.

106. Lan Y, He L, Dong X, Tang R, Li W, Wang J, Wang L, Yue B, Price M, Guo T et al: Comparative transcriptomes of three different skin sites for the Asiatic toad *B*(*ufo gargarizans*). PeerJ 2022, 10:e12993.

107. Lv Y, Li Y, Wen Z, Shi Q: Transcriptomic and gene-family dynamic analyses reveal gene expression pattern and evolution in toxin-producing tissues of Asiatic toad (*Bufo gargarizans*). Frontiers in Ecology and Evolution 2022, 10:924248.

108. Mohammadi S, Herrera-Álvarez S, Yang L, Rodriguez-Ordonez MdP, Zhang K, Storz JF, Dobler S, Crawford AJ, Andolfatto P: Constraints on the evolution of toxin-resistant Na, K-ATPases have limited dependence on sequence divergence. PLoS Genet 2022, 18(8):e1010323.

109. Mohammadi S, Yang L, Harpak A, Herrera-Álvarez S, del Pilar Rodríguez-Ordoñez M, Peng J, Zhang K, Storz JF, Dobler S, Crawford AJ: Concerted evolution reveals co-adapted amino acid substitutions in Na+ K+-ATPase of frogs that prey on toxic toads. Curr Biol 2021, 31(12):2530–2538. e2510.

110. Hutchinson DA, Mori A, Savitzky AH, Burghardt GM, Wu X, Meinwald J, Schroeder FC: Dietary sequestration of defensive steroids in nuchal glands of the Asian snake *Rhabdophis tigrinus*. Proceedings of the National Academy of Sciences 2007, 104(7):2265–2270.

111. Brodie III ED, Brodie Jr ED: Tetrodotoxin resistance in garter snakes: an evolutionary response of predators to dangerous prey. Evolution 1990, 44(3):651–659.

112. Mancuso M, Zaman S, Maddock ST, Kamei RG, Salazar-Valenzuela D, Wilkinson M, Roelants K, Fry BG: Resistance Is Not Futile: Widespread Convergent Evolution of Resistance to Alpha-Neurotoxic Snake Venoms in Caecilians (Amphibia: Gymnophiona). International Journal of Molecular Sciences 2023, 24(14):11353.

113. Symula R, Schulte R, Summers K: Molecular phylogenetic evidence for a mimetic radiation in Peruvian poison frogs supports a Müllerian mimicry hypothesis. Proceedings of the Royal Society of London Series B: Biological Sciences 2001, 268(1484):2415–2421.

114. Daly JW, Brown GB, Mensah-Dwumah M, Myers CW: Classification of skin alkaloids from neotropical poison-dart frogs (Dendrobatidae). Toxicon 1978, 16(2):163–188.

115. Stuckert AMM, Moore E, Coyle KP, Davison I, MacManes MD, Roberts R, Summers K: Variation in pigmentation gene expression is associated with distinct aposematic color morphs in the poison frog *Dendrobates auratus*. BMC Evol Biol 2019, 19(1):85.

116. Stuckert AM, Chouteau M, McClure M, LaPolice TM, Linderoth T, Nielsen R, Summers K, MacManes MD: The genomics of mimicry: gene expression throughout development provides insights into convergent and divergent phenotypes in a Müllerian mimicry system. Mol Ecol 2021, 30(16):4039–4061.

117. Twomey E, Johnson JD, Castroviejo-Fisher S, Van Bocxlaer I: A ketocarotenoid-based colour polymorphism in the Sira poison frog *Ranitomeya sirensis* indicates novel gene interactions underlying aposematic signal variation. Mol Ecol 2020, 29(11):2004–2015.

118. Twomey E, Kain M, Claeys M, Summers K, Castroviejo-Fisher S, Van Bocxlaer I: Mechanisms for color convergence in a mimetic radiation of poison frogs. The American Naturalist 2020, 195(5):E132–E149.

119. Linderoth T, Aguilar-Gómez D, White E, Twomey E, Stuckert A, Bi K, Ko A, Graham N, Rocha JL, Chang J et al: Genetic basis of aposematic coloration in a mimetic radiation of poison frogs. bioRxiv 2023:2023.2004.2020.537757.

120. Stuckert AM, Freeborn L, Howell KA, Yang Y, Nielsen R, Richards-Zawacki C, MacManes MD: Transcriptomic analyses during development reveal mechanisms of integument structuring and color production. Evol Ecol 2023:1–22.

121. Burgon JD, Vieites DR, Jacobs A, Weidt SK, Gunter HM, Steinfartz S, Burgess K, Mable BK, Elmer KR: Functional colour genes and signals of selection in colour-polymorphic salamanders. Mol Ecol 2020, 29(7):1284–1299.

122. Fischer EK, Roland AB, Moskowitz NA, Tapia EE, Summers K, Coloma LA, O’Connell LA: The neural basis of tadpole transport in poison frogs. Proceedings of the Royal Society B: Biological Sciences 2019, 286(1907):20191084.

123. Crump ML: Parental Care among the Amphibia. In: Advances in the Study of Behavior. Edited by Rosenblatt JS, Snowdon CT, vol. 25: Academic Press; 1996: 109–144.

124. Mailho-Fontana PL, Antoniazzi MM, Coelho GR, Pimenta DC, Fernandes LP, Kupfer A, Brodie ED, Jared C: Milk provisioning in oviparous caecilian amphibians. Science 2024, 383(6687):1092–1095.

125. Kupfer A, Müller H, Antoniazzi MM, Jared C, Greven H, Nussbaum RA, Wilkinson M: Parental investment by skin feeding in a caecilian amphibian. Nature 2006, 440(7086):926–929.

126. Liu Y, Jones CD, Day LB, Summers K, Burmeister SS: Cognitive phenotype and differential gene expression in a hippocampal homologue in two species of frog. Integr Comp Biol 2020, 60(4):1007–1023.

127. Wu W, Gao YD, Jiang DC, Lei J, Ren JL, Liao WB, Deng C, Wang Z, Hillis DM, Zhang YP et al: Genomic adaptations for arboreal locomotion in Asian flying treefrogs. Proc Natl Acad Sci U S A 2022, 119(13):e2116342119.

128. Blackburn DC, Gray JA, Stanley EL: The only “lungless” frog has a glottis and lungs. Curr Biol 2024, 34(10):R492–R493.

129. Lewis ZR, Kerney R, Hanken J: Developmental basis of evolutionary lung loss in plethodontid salamanders. Sci Adv 2022, 8(33):eabo6108.

130. Heiss E, Natchev N, Salaberger D, Gumpenberger M, Rabanser A, Weisgram J: Hurt yourself to hurt your enemy: new insights on the function of the bizarre antipredator mechanism in the salamandrid *Pleurodeles waltl*. J Zool 2010, 280(2):156–162.

131. Brodie Jr ED, Nussbaum RA, DiGiovanni M: Antipredator adaptations of Asian salamanders (Salamandridae). Herpetologica 1984:56–68.

132. Luedtke JA, Chanson J, Neam K, Hobin L, Maciel AO, Catenazzi A, Borzée A, Hamidy A, Aowphol A, Jean A et al: Ongoing declines for the world’s amphibians in the face of emerging threats. Nature 2023, 622(7982):308–314.

133. Vacher JP, Chave J, Ficetola FG, Sommeria-Klein G, Tao S, Thébaud C, Blanc M, Camacho A, Cassimiro J, Colston TJ: Large-scale DNA-based survey of frogs in Amazonia suggests a vast underestimation of species richness and endemism. J Biogeogr 2020, 47(8):1781–1791.

134. Oliver PM, Bower DS, McDonald PJ, Kraus F, Luedtke J, Neam K, Hobin L, Chauvenet AL, Allison A, Arida E: Melanesia holds the world’s most diverse and intact insular amphibian fauna. Communications biology 2022, 5(1):1182.

135. Liu J, Slik F, Zheng S, Lindenmayer DB: Undescribed species have higher extinction risk than known species. Conservation Letters 2022, 15(3):e12876.

136. Re:wild, Earth S, Group ISAS: State of the World’s Amphibians: The Second Global Amphibian Assessment. In. Texas, USA: Re:wild; 2023.

137. Gower DJ, San Mauro D, Giri V, Bhatta G, Govindappa V, Kotharambath R, Oommen OV, Fatih FA, Mackenzie-Dodds JA, Nussbaum RA: Molecular systematics of caeciliid caecilians (Amphibia: Gymnophiona) of the Western Ghats, India. Mol Phylogen Evol 2011, 59(3):698–707.

138. Forester BR, Beever EA, Darst C, Szymanski J, Funk WC: Linking evolutionary potential to extinction risk: applications and future directions. Front Ecol Environ 2022, 20(9):507–515.

139. Zhang Y, Stern AJ, Nielsen R: The evolutionary dynamics of local adaptations under genetic rescue is determined by mutational load and polygenicity. J Hered 2024, 115(4):373–384.

140. Nemesházi E, Bókony V: Interplay of genotypic and thermal sex determination shapes climatic distribution in herpetofauna. bioRxiv 2024:2024.2004.2021.589911.

141. Wollenberg Valero KC, Marshall JC, Bastiaans E, Caccone A, Camargo A, Morando M, Niemiller ML, Pabijan M, Russello MA, Sinervo B et al: Patterns, Mechanisms and Genetics of Speciation in Reptiles and Amphibians. Genes (Basel)2019, 10(9):646.

142. Wren S, Borzee A, Marcec-Greaves R, Angulo A: Amphibian conservation action plan : a status review and roadmap for global amphibian conservation. Gland, Switzerland : IUCN, 2024: IUCN; 2024.

143. Hogg CJ: Translating genomic advances into biodiversity conservation. Nat Rev Genet 2023, 25(5):362–373.

144. Whiteley AR, Fitzpatrick SW, Funk WC, Tallmon DA: Genetic rescue to the rescue. Trends Ecol Evol 2015, 30(1):42–49.

145. Pabijan M, Palomar G, Antunes B, Antoł W, Zieliński P, Babik W: Evolutionary principles guiding amphibian conservation. Evol Appl 2020, 13(5):857–878.

146. Theissinger K, Fernandes C, Formenti G, Bista I, Berg PR, Bleidorn C, Bombarely A, Crottini A, Gallo GR, Godoy JA et al: How genomics can help biodiversity conservation. Trends Genet 2023, 39(7):545–559.

147. Trumbo D, Hardy B, Crockett H, Muths E, Forester B, Cheek R, Zimmerman S, Corey-Rivas S, Bailey L, Funk WC: Conservation genomics of an endangered montane amphibian reveals low population structure, low genomic diversity, and selection pressure from disease. Mol Ecol 2023, 32(24):6777–6795.

148. Torres-Sánchez M, Longo AV: Linking pathogen-microbiome-host interactions to explain amphibian population dynamics. Mol Ecol 2022, 31(22):5784–5794.

149. Allendorf FW, Funk WC, Aitken SN, Byrne M, Luikart G: Conservation and the genomics of populations: Oxford University Press; 2022.

150. Fischman RL, Ruhl JB, Forester BR, Lama TM, Kardos M, Rojas GA, Robinson NA, Shirey PD, Lamberti GA, Ando AW et al: A landmark environmental law looks ahead. Science 2023, 382(6677):1348–1355.

151. Programme UE: Kunming-Montreal Global Biodiversity Framework — CBD/COP/15/L25. In. Edited by Diversity UCoB. Montreal Canada; 2022.

152. Formenti G, Theissinger K, Fernandes C, Bista I, Bombarely A, Bleidorn C, Ciofi C, Crottini A, Godoy JA, Höglund J et al: The era of reference genomes in conservation genomics. Trends Ecol Evol 2022.

153. Kosch TA, Waddle AW, Cooper CA, Zenger KR, Garrick DJ, Berger L, Skerratt LF: Genetic approaches for increasing fitness in endangered species. Trends Ecol Evol 2022, 37(4):332–345.

154. Kosch T, Silva C, Brannelly L, Roberts A, Lau Q, Marantelli G, Berger L, Skerratt L: Genetic potential for disease resistance in critically endangered amphibians decimated by chytridiomycosis. Anim Conserv 2019, 22(3):238–250.

155. Savage AE, Gratwicke B, Hope K, Bronikowski E, Fleischer RC: Sustained immune activation is associated with susceptibility to the amphibian chytrid fungus. Mol Ecol 2020, 29(15):2889–2903.

156. Bataille A, Cashins SD, Grogan L, Skerratt LF, Hunter D, McFadden M, Scheele B, Brannelly LA, Macris A, Harlow PS: Susceptibility of amphibians to chytridiomycosis is associated with MHC class II conformation. Proc R Soc Lond, Ser B: Biol Sci 2015, 282(1805):20143127.

157. Knapp RA, Wilber MQ, Byrne AQ, Joseph MB, Smith TC, Rothstein AP, Grasso RL, Rosenblum EB: Reintroduction of resistant frogs facilitates landscape-scale recovery in the presence of a lethal fungal disease. bioRxiv 2023:2023.2005.2022.541534.

158. Byrne PG, Silla AJ: An experimental test of the genetic consequences of population augmentation in an amphibian. Conserv Sci Pract 2020, 2(6):e194.

159. Liddell E, Sunnucks P, Cook CN: To mix or not to mix gene pools for threatened species management? Few studies use genetic data to examine the risks of both actions, but failing to do so leads disproportionately to recommendations for separate management. Biol Conserv 2021, 256:109072.

160. Kyriazis CC, Wayne RK, Lohmueller KE: Strongly deleterious mutations are a primary determinant of extinction risk due to inbreeding depression. Evolution Letters 2021, 5:33–47.

161. Berger L, Skerratt LF, Kosch TA, Brannelly LA, Webb RJ, Waddle AW: Advances in Managing Chytridiomycosis for Australian Frogs:*Gradarius Firmus* Victoria. Annu Rev Anim Biosci 2024, 12(1):113–133.

162. Wong K-C: Big data challenges in genome informatics. Biophysical Reviews 2019, 11(1):51–54.

163. Zuo B, Nneji LM, Sun Y-B: Comparative genomics reveals insights into anuran genome size evolution. BMC Genomics 2023, 24(1):379.

164. Treangen TJ, Salzberg SL: Repetitive DNA and next-generation sequencing: computational challenges and solutions. Nat Rev Gene*t* 2012, 13(1):36–46.

165. Alkan C, Sajjadian S, Eichler EE: Limitations of next-generation genome sequence assembly. Nat Methods 2011, 8(1):61–65.

166. Mable B, Alexandrou M, Taylor M: Genome duplication in amphibians and fish: an extended synthesis. J Zool 2011, 284(3):151–182.

167. Schmid M, Evans BJ, Bogart JP: Polyploidy in Amphibia. Cytogenet Genome Res 2015, 145(3-4):315–330.

168. Sun Y, Shang L, Zhu Q-H, Fan L, Guo L: Twenty years of plant genome sequencing: achievements and challenges. Trends Plant Sci 2022, 27(4):391–401.

169. Session AM, Uno Y, Kwon T, Chapman JA, Toyoda A, Takahashi S, Fukui A, Hikosaka A, Suzuki A, Kondo M et al: Genome evolution in the allotetraploid frog*Xenopus laevis*. Nature 2016, 538(7625):336–343.

170. Rhie A, McCarthy SA, Fedrigo O, Damas J, Formenti G, Koren S, Uliano-Silva M, Chow W, Fungtammasan A, Kim J et al: Towards complete and error-free genome assemblies of all vertebrate species. Nature 2021, 592(7856):737–746.

171. Consortium TU: UniProt: the Universal Protein Knowledgebase in 2023. Nucleic Acids Res 2022, 51(D1):D523–D531.

172. Dimitrakopoulou D, Khwatenge CN, James-Zorn C, Paiola M, Bellin EW, Tian Y, Sundararaj N, Polak EJ, Grayfer L, Barnard D et al: Advances in the *Xenopus* immunome: Diversification, expansion, and contraction. Dev Comp Immunol 2023, 145:104734.

173. Silla AJ, Byrne PG: The Role of Reproductive Technologies in Amphibian Conservation Breeding Programs. Annu Rev Anim Biosci 2019, 7(1):499–519.

174. Banach M, Edholm E-S, Robert J: Exploring the functions of nonclassical MHC class Ib genes in *Xenopus laevis* by the CRISPR/Cas9 system. Dev Biol 2016, 426(2):261–269.

175. Grainger RM: *Xenopus tropicalis* as a model organism for genetics and genomics: past, present, and future. Xenopus Protocols: Post-Genomic Approaches 2012:3–15.

176. Sousounis K, Courtemanche K, Whited JL: A Practical Guide for CRISPR-Cas9-Induced Mutations in Axolotls. In: Salamanders: Methods and Protocols. Springer; 2022: 335–349.

177. Wang F, Shi Z, Cui Y, Guo X, Shi Y-B, Chen Y: Targeted gene disruption in*Xenopus laevis* using CRISPR/Cas9. Cell & bioscience 2015, 5:1–5.

178. Douglas AJ, Todd LA, Katzenback BA: The amphibian invitrome: Past, present, and future contributions to our understanding of amphibian immunity. Dev Comp Immunol 2023, 142:104644.

179. Bui-Marinos MP, Todd LA, Douglas AJ, Katzenback BA: So, you want to create a frog cell line? A guide to establishing frog skin cell lines from tissue explants. MethodsX 2022, 9:101693.

180. Dahn HA, Mountcastle J, Balacco J, Winkler S, Bista I, Schmitt AD, Pettersson OV, Formenti G, Oliver K, Smith M et al: Benchmarking ultra-high molecular weight DNA preservation methods for long-read and long-range sequencing. GigaScience 2022, 11.

181. Forzán MJ, Heatley J, Russell KE, Horney B: Clinical pathology of amphibians: a review. Veterinary Clinical Pathology 2017, 46(1):11–33.

182. IUCN: IUCN policy statement on research involving species at risk of extinction. In. Gland, Switzerland: International Union for the Conservation of Nature.; 1989.

183. Ambu J, Dufresnes C: Buccal swabs for amphibian genomics. Amphibia-Reptilia 2023, 44(2):249–255.

184. Oyler-McCance SJ, Ryan MJ, Sullivan BK, Fike JA, Cornman RS, Giermakowski JT, Zimmerman SJ, Harrow RL, Hedwall SJ, Hossack BR et al: Genetic connectivity in the Arizona toad (*Anaxyrus microscaphus*): implications for conservation of a stream dwelling amphibian in the arid Southwestern United States. Conserv Genet 2024, 25(3):835–848.

185. Farquharson KA, McLennan EA, Belov K, Hogg CJ: The genome sequence of the critically endangered Kroombit tinkerfrog (*Taudactylus pleione*). F1000Research 2023, 12:845.

186. Fong JJ, Blom MP, Aowphol A, McGuire JA, Sutcharit C, Soltis PS: Recent advances in museomics: revolutionizing biodiversity research. Frontiers in Ecology and Evolution 2023, 11:1188172.

187. Scherz MD, Rasolonjatovo SM, Köhler J, Rancilhac L, Rakotoarison A, Raselimanana AP, Ohler A, Preick M, Hofreiter M, Glaw F: ‘Barcode fishing’ for archival DNA from historical type material overcomes taxonomic hurdles, enabling the description of a new frog species. Sci Rep 2020, 10(1):19109.

188. Rancilhac L, Bruy T, Scherz MD, Pereira EA, Preick M, Straube N, Lyra ML, Ohler A, Streicher JW, Andreone F: Target-enriched DNA sequencing from historical type material enables a partial revision of the Madagascar giant stream frogs (genus *Mantidactylus*). J Nat Hist 2020, 54(1-4):87–118.

189. Rakotoarison A, Scherz MD, Mullin KE, Crottini A, Petzold A, Ranjanaharisoa FA, Maheritafika HMR, Rafanoharana JM, Raherinjatovo H, Andreone F: Gray versus yellow ventral coloration: Identity, distribution, color polymorphism and molecular relationships of the microhylid frog *Platypelis mavomavo* Andreone, Fenolio & Walvoord, 2003. Zootaxa 2023, 5352(2):221–234.

190. Evans BJ, Gansauge M-T, Stanley EL, Furman BLS, Cauret CMS, Ofori-Boateng C, Gvoždík V, Streicher JW, Greenbaum E, Tinsley RC et al: *Xenopus fraseri* : Mr. Fraser, where did your frog come from? PLOS ONE 2019, 14(9):e0220892.

191. Raxworthy CJ, Smith BT: Mining museums for historical DNA: advances and challenges in museomics. Trends Ecol Evol 2021, 36(11):1049–1060.

192. Roycroft E, Moritz C, Rowe KC, Moussalli A, Eldridge MDB, Portela Miguez R, Piggott MP, Potter S: Sequence Capture From Historical Museum Specimens: Maximizing Value for Population and Phylogenomic Studies. Frontiers in Ecology and Evolution 2022, 10:931644.

193. Speer KA, Hawkins MTR, Flores MFC, McGowen MR, Fleischer RC, Maldonado JE, Campana MG, Muletz-Wolz CR: A comparative study of RNA yields from museum specimens, including an optimized protocol for extracting RNA from formalin-fixed specimens. Frontiers in Ecology and Evolution 2022, 10:953131.

194. Dabney J, Knapp M, Glocke I, Gansauge M-T, Weihmann A, Nickel B, Valdiosera C, García N, Pääbo S, Arsuaga J-L: Complete mitochondrial genome sequence of a Middle Pleistocene cave bear reconstructed from ultrashort DNA fragments. Proceedings of the National Academy of Sciences 2013, 110(39):15758–15763.

195. Gansauge M-T, Gerber T, Glocke I, Korlević P, Lippik L, Nagel S, Riehl LM, Schmidt A, Meyer M: Single-stranded DNA library preparation from highly degraded DNA using T4 DNA ligase. Nucleic Acids Res 2017, 45(10):e79–e79.

196. Straube N, Lyra ML, Paijmans JL, Preick M, Basler N, Penner J, Rödel MO, Westbury MV, Haddad CF, Barlow A: Successful application of ancient DNA extraction and library construction protocols to museum wet collection specimens. Mol Ecol Resour 2021, 21(7):2299–2315.

197. Dalén L, Heintzman PD, Kapp JD, Shapiro B: Deep-time paleogenomics and the limits of DNA survival. Science 2023, 382(6666):48–53.

198. Ramírez JP, Jaramillo CA, Lindquist ED, Crawford AJ, Ibáñez R: Recent and Rapid Radiation of the Highly Endangered Harlequin Frogs (*Atelopus*) into Central America Inferred from Mitochondrial DNA Sequences. Diversity 2020, 12(9):360.

199. Hutter CR, Cobb KA, Portik DM, Travers SL, Wood Jr. PL, Brown RM: FrogCap: A modular sequence capture probe-set for phylogenomics and population genetics for all frogs, assessed across multiple phylogenetic scales. Mol Ecol Resour 2022, 22(3):1100–1119.

200. Renner SS, Scherz MD, Schoch CL, Gottschling M, Vences M: DNA sequences from type specimens and type strains–how to increase their number and improve their annotation in NCBI GenBank and related databases. Syst Biol 2023:syad068.

201. Alves RJV, Weksler M, Oliveira JA, Buckup PA, Santana HR, Peracchi AL, Kellner AW, Aleixo A, Langguth A, Almeida A: Brazilian legislation on genetic heritage harms Biodiversity Convention goals and threatens basic biology research and education. In., vol. 90: SciELO Brasil; 2018: 1279–1284.

202. Alexander GJ, Tollev KA, Maritz B, McKechnie A, Manger P, Thomson RL, Schradin C, Fuller A, Meyer L, Hetem RS: Excessive red tape is strangling biodiversity research in South Africa. S Afr J Sci 2021, 117(9-10):1–4.

203. Collier-Robinson L, Rayne A, Rupene M, Thoms C, Steeves T: Embedding indigenous principles in genomic research of culturally significant species. N Z J Ecol 2019, 43(3):1–9.

204. Rayne A, Blair S, Dale M, Flack B, Hollows J, Moraga R, Parata RN, Rupene M, Tamati-Elliffe P, Wehi PM et al: Weaving place-based knowledge for culturally significant species in the age of genomics: Looking to the past to navigate the future. Evol Appl 2022, 15(5):751–772.

205. Mc Cartney AM, Head MA, Tsosie KS, Sterner B, Glass JR, Paez S, Geary J, Hudson M: Indigenous peoples and local communities as partners in the sequencing of global eukaryotic biodiversity. npj Biodiversity 2023, 2(1):8.

206. Carroll S, Garba I, Figueroa-Rodríguez O, Holbrook J, Lovett R, Materechera S, Parsons M, Raseroka K, Rodriguez-Lonebear D, Rowe R: The CARE principles for indigenous data governance. Data Science Journal 2020, 19(43):1–21.

207. Ramírez-Castañeda V, Westeen EP, Frederick J, Amini S, Wait DR, Achmadi AS, Andayani N, Arida E, Arifin U, Bernal MA et al: A set of principles and practical suggestions for equitable fieldwork in biology. Proceedings of the National Academy of Sciences 2022, 119(34):e2122667119.

208. Lewin HA, Richards S, Lieberman Aiden E, Allende ML, Archibald JM, Bálint M, Barker KB, Baumgartner B, Belov K, Bertorelle G et al: The Earth BioGenome Project 2020: Starting the clock. Proceedings of the National Academy of Sciences 2022, 119(4):e2115635118.

209. Stiller J, Feng S, Chowdhury A-A, Rivas-González I, Duchêne DA, Fang Q, Deng Y, Kozlov A, Stamatakis A, Claramunt S et al: Complexity of avian evolution revealed by family-level genomes. Nature 2024, 629(818):851–860.

210. Feres MVC: Biodiversity, traditional knowledge and patent rights: The case study of *Phyllomedusa bicolor*. Revista Direito GV 2022, 18:e2205.

211. Te Aika B, Liggins L, Rye C, Perkins EO, Huh J, Brauning R, Godfery T, Black MA: Aotearoa genomic data repository: An āhuru mōwai for taonga species sequencing data. Mol Ecol Resour 2023, 00:1–14.

212. Buckner JC, Sanders RC, Faircloth BC, Chakrabarty P: The critical importance of vouchers in genomics. eLife 2021, 10:e68264.

213. Golan J, Riddle K, Hudson M, Anderson J, Kusabs N, Coltman T: Benefit sharing: Why inclusive provenance metadata matter. Front Genet 2022, 13:1014044.

214. Haelewaters D, Hofmann TA, Romero-Olivares AL: Ten simple rules for Global North researchers to stop perpetuating helicopter research in the Global South. PLoS Comp Biol 2021, 17(8):e1009277.

215. de Vos A, Schwartz MW: Confronting parachute science in conservation. Conserv Sci Pract 2022, 4(5):e12681.

216. Watsa M, Erkenswick GA, Pomerantz A, Prost S: Portable sequencing as a teaching tool in conservation and biodiversity research. PLoS Biol 2020, 18(4):e3000667.

217. Armenteras D: Guidelines for healthy global scientific collaborations. Nature Ecology & Evolution 2021, 5(9):1193–1194.

