## Supplemental information for "The Amphibian Genomics Consortium: advancing genomic and genetic resources for amphibian research and conservation"

### This PDF includes:

1. Supplementary Methods

2. Table S1

3. Figures S1 – S3

4. References

**Supplementary Methods**

We downloaded records from the NCBI Genome browser to compare the number of sequenced genomes and genome size across tetrapod classes (all NCBI records were accessed on 1 March 2024). Current number of species per class was obtained from different databases (amphibians: <https://amphibiaweb.org/>, reptiles: <http://www.reptile-database.org/>, birds: <https://datazone.birdlife.org/>, mammals: <https://www.mammaldiversity.org/>; records accessed on 1 March 2024). We built a contingency table with the numbers of total species and the number of species with sequenced genomes for each tetrapod class, compared these values through a Chi-squared test with the *chisq.test* function of R, and visualized data percentages using a mosaic plot by the *mosaicplot* function of R. To assess genome size bias between species with and without sequenced genomes across tetrapods, we also downloaded genome size records from the Animal Genome Size Database ([https://www.genomesize.com/ 1]). We averaged genome size C-values for species with more than one estimate and calculated genome size in Gb considering 1 pg equivalent to 0.978 Gb. Due to the low number of amphibian records for caecilians, we obtained additional amphibian genome size values from HC Liedtke, DJ Gower, M Wilkinson and I Gomez-Mestre [2]. The raincloud plots visualized data from NCBI genome assembly sizes and estimated genome size records of unsequenced species. The plots used combined box and density plots with raw values as points (<https://github.com/PsyTeachR/introdataviz>). We used this graphical representation for exploring genome differences across tetrapod classes and amphibian orders, respectively. We formally tested differences in genome size by tetrapod class for species with and without reference genomes using analysis of the variance and post-hoc tests with the *aov* and *TukeyHSD* functions of R. We further explored the distribution of genome sizes regarding genome references at family level in the amphibian phylogeny from W Jetz and RA Pyron [3] including genome size inferences from GoaT [4] using *phytools* package of R [5]. We also summarized amphibian information from NCBI Sequence Read Archive (SRA) by sorting amphibian SRA data by BioProject ID and quantifying the number of times different sequencing techniques have been applied to the study of amphibians. We represented this information by grouping BioProjects yearly.

To analyze the information from the results of the AGC survey (see Table S2 for detailed information about AGC survey questions), we first quantified the number of sequencing techniques that AGC members have used in their research and the amphibian order that these researchers have worked with (AGC survey questions 11 and 9). We visualized both datasets using bar plots with *barplot* function of R. Answers from questions 8, 12, and 13 treated as quantitative variables were combined using principal components analysis with the *prcomp* function of R. We visualized the results of this dimensionality reduction analysis using several plots with the functions of *factorextra* package of R [6]. We created two new variables with the scores of the first and second dimensions, which grouped answers defining genomics expertise and identified challenges of the AGC members. To explain the variation of these two new variables, we used as explanatory variables the scientific expertise of AGC members (AGC survey question 5), the funding success (AGC survey question 16), and two other variables based on the information provided about the country of main affiliation of the member (AGC survey question 4). These two variables were the number of described amphibian species (data from AmphibiaWeb: <https://amphibiaweb.org/>) and gross domestic expenditure on R&D (GERD) per capita (data from UNESCO: http://data.uis.unesco.org, World Bank: https://data.worldbank.org/, and World population data: <https://worldpopulationreview.com/>). We explored the correlation between the dependent (scores from dimension 1 and 2) and independent variables and collinearity among independent variables and, subsequently, modelled dimension 1 and dimension 2 independently using generalized linear models with *glm* function of R.

**Table S1. Amphibian Genomics Consortium (AGC) Survey Questions.** Each row contains one of the 23 questions.

**1** Contact details (answering this question is optional; all data will be treated anonymously and details will only be used to contact respondents if clarifications on answers are required)

Name	(1) __________________________________________________

E-mail	(2) __________________________________________________

**2** Are you member of the Amphibian Genomics Consortium (https://mvs.unimelb.edu.au/amphibian-genomics-consortium)?

Yes (1)

No (2)

**3** Organization(s) you are affiliated to (write the name of your organization in the corresponding category)

University	(1) _____________________________________________

Research center / Museum	(2) _____________________________________________

Private company	(3) _____________________________________________

NGO	(4) _____________________________________________

**4** Country(ies) of affiliation

__________________________________________________

**5** Number of years since PhD dissertation, as a measure of research experience. Indicate “0” if you are a student (i.e., Master’s, PhD student/candidate, or similar)

__________________________________________________

**6** Research topic. Rate each topic, being 0 = you haven’t worked in the field yet, 1 = rarely worked, 2 = occasionally, 3 = often, and 4 = main research topicç

Ecology and Evolution	(1) _________________________________

Conservation and Management	(2) _________________________________

Development and Regeneration	(3) _________________________________

Biomedical research and drug discovery	(4) _________________________________

**7** How often have you applied sequencing technologies in your research? Indicate a number, being 0 = never, 1 = rarely, 2 = occasionally, 3 = often, and 4 = mainly

__________________________________________________

**8** Type of work you mainly undertake. Rate each of the following items, being 0 = you haven’t performed the task yet, 1 = rarely performed, 2 = occasionally, 3 = often, and 4 = main activity

Field surveys	(1) _________________________________

Laboratory work	(2) _________________________________

Computational work	(3) _________________________________

Writing (articles, reports, and grants)	(4) _________________________________

**9** Amphibian order you mainly work with. Rate each of the amphibian orders, being 0 = you haven’t worked with the group yet, 1 = rarely worked with it, 2 = occasionally, 3 = often, and 4 = mainly

Anura	(1) __________________________________________________

Caudata	(2) __________________________________________________

Gymnophiona	(3) __________________________________________________

**10** Select sequencing technologies that you have implemented in your research (multiple choice)

Sanger sequencing (1)

Short read sequencing (2)

Long read sequencing (3)

**11** Select the sequencing focus of your research (multiple choice)

Whole genome sequencing (1)

Whole exome sequencing (2)

Whole methylome sequencing (3)

Chromatin Immunoprecipitation followed by sequencing (4)

Transcriptome sequencing (5)

Restriction site-associated DNA sequencing (6)

Targeted sequencing (7)

Other (8)

**12** Bioinformatic analyses that you have performed. Rate each of the following analyses by your level of expertise, being 0 = you haven’t performed the analysis yet, 1 = novice at the analysis, 2 = advanced beginner, 3 = competent, 4 = proficient, and 5 = expert

Sequences quality control	(1) _____________________________________________

Filtering and trimming	(2) _____________________________________________

De novo assembly	(3) _____________________________________________

Mapping to reference	(4) _____________________________________________

Genomics annotation	(5) _____________________________________________

Downstream analyses	(6) _____________________________________________

**13** Evaluate the challenges for the amphibian genomics advancement. Rate each of the following items, being 0 = not issue at all, 1 = somehow problematic, 2 = average, 3 = complicated, and 4 = extremely challenging

Sample acquisition	(1) _____________________________________________

Protocol transferability	(2) _____________________________________________

Data quantity and quality	(3) _____________________________________________

Data integration	(4) _____________________________________________

Computational resources	(5) _____________________________________________

Data storage	(6) _____________________________________________

Funding sources	(7) _____________________________________________

**14** Identify other challenges and score them according to the scale in the previous question

__________________________________________________

**15** Have you applied for any funding for high-throughput sequencing in amphibians?

Yes (1)

No (2)

**16** If yes in question 15, have you been successful?

Yes (1)

No (2)

**17** Type of body fund/funds that you have applied (multiple choice)

Governmental (1)

Scientific association (2)

Private company (3)

NGO (4)

Other (5)

**18** Name the body fund/funds

__________________________________________________

**19** How much money have you roughly obtained specifically for sequencing (for large projects please consider only the subtotal amount related to sample preparation and sequencing)

0 (1)

0 – 5,000 (2)

5,000 – 10,000 (3)

10,000 – 50,000 (4)

>50,000 (5)

**20** Money currency of the sequencing project(s)

__________________________________________________

**21** Type of body fund(s) that could support amphibian genomics initiatives, such as AGC actions (multiple choice)

Governmental (1)

Scientific association (2)

Private company (3)

NGO (4)

Other (5)

**22** Name the body fund(s)s that could support amphibian genomics initiatives, such as AGC actions

__________________________________________________

**23** State in a sentence or two your opinion about the current gap of knowledge in amphibian genomics

__________________________________________________

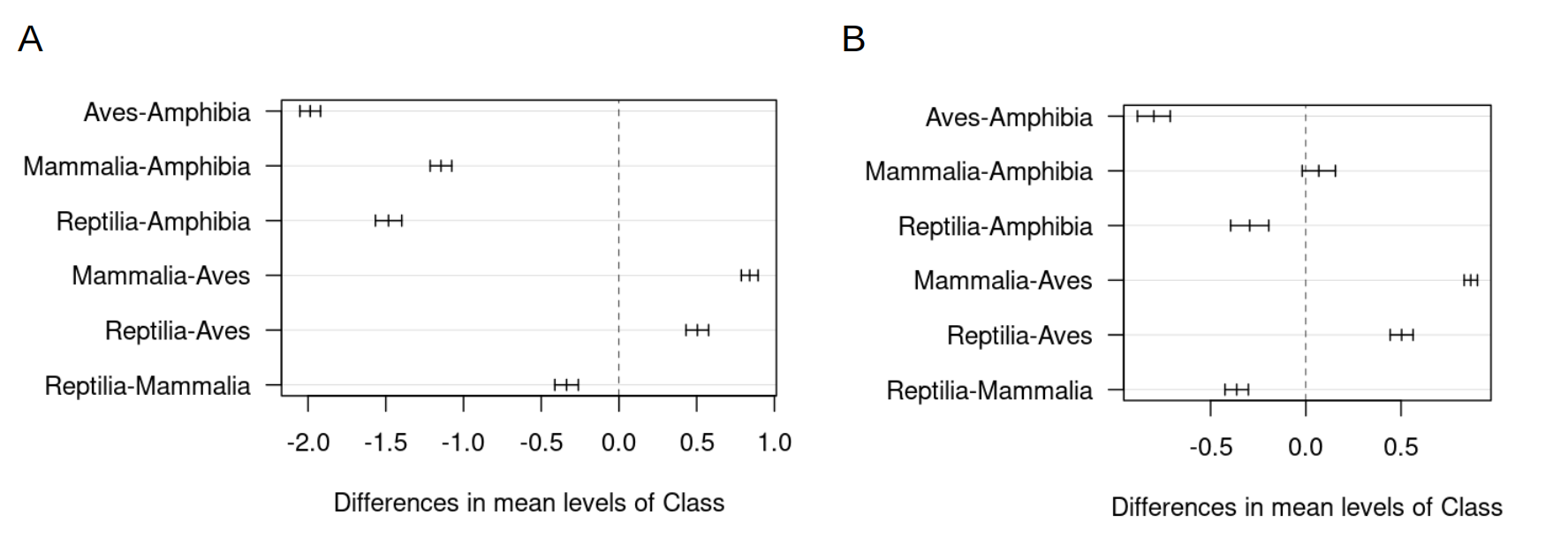

**Figure S1. Mean differences of genome size across tetrapods.** Comparisons of all possible pairs of tetrapod classes resulted from the post-hoc analyses using Tukey tests. Charts represent the mean differences for each pair of class (e.g. Aves-Amphibia) using all the species explored in our study (A) and only the species sequenced genomes (B).

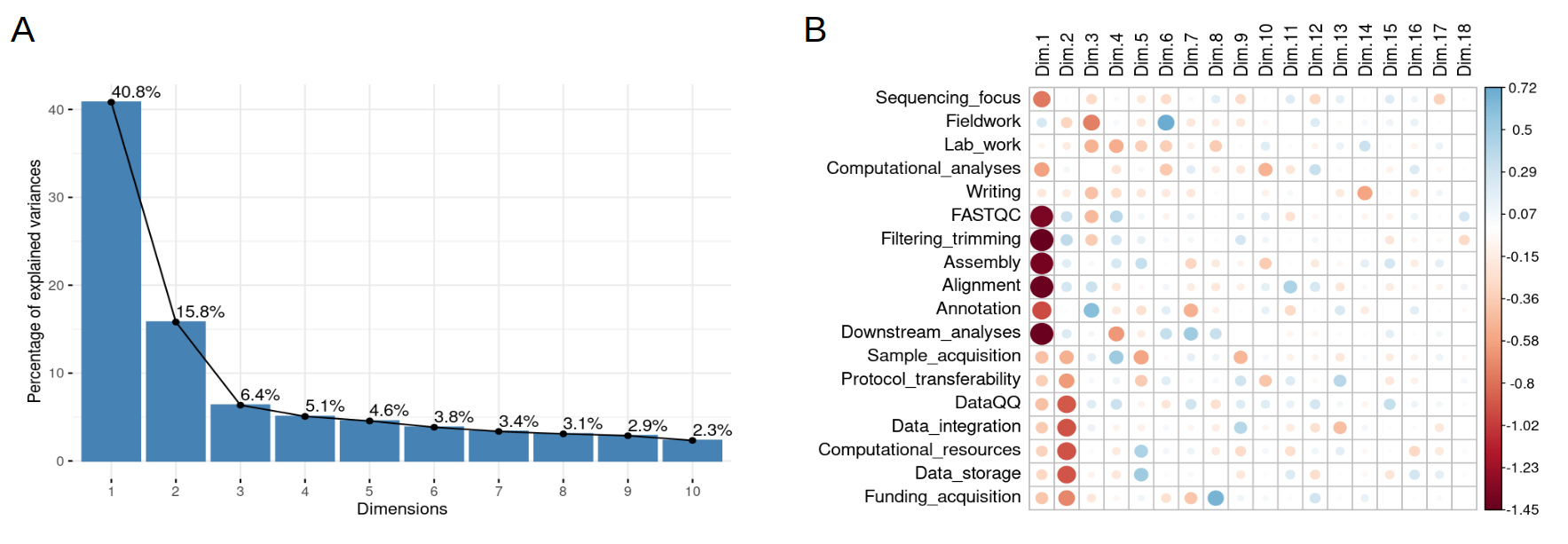

**Figure S2. Principal component analysis (PCA) results of the responses of the Amphibian Genomics Consortium (AGC) survey**. (A) Bar plot representing the percentage of explained variance per each dimension. (B) Correlation of each variable (from the quantitative questions of the Amphibian Genomics Consortium [AGC] survey) with each PCA dimension. Direction of the relation and effect is color and size coded.

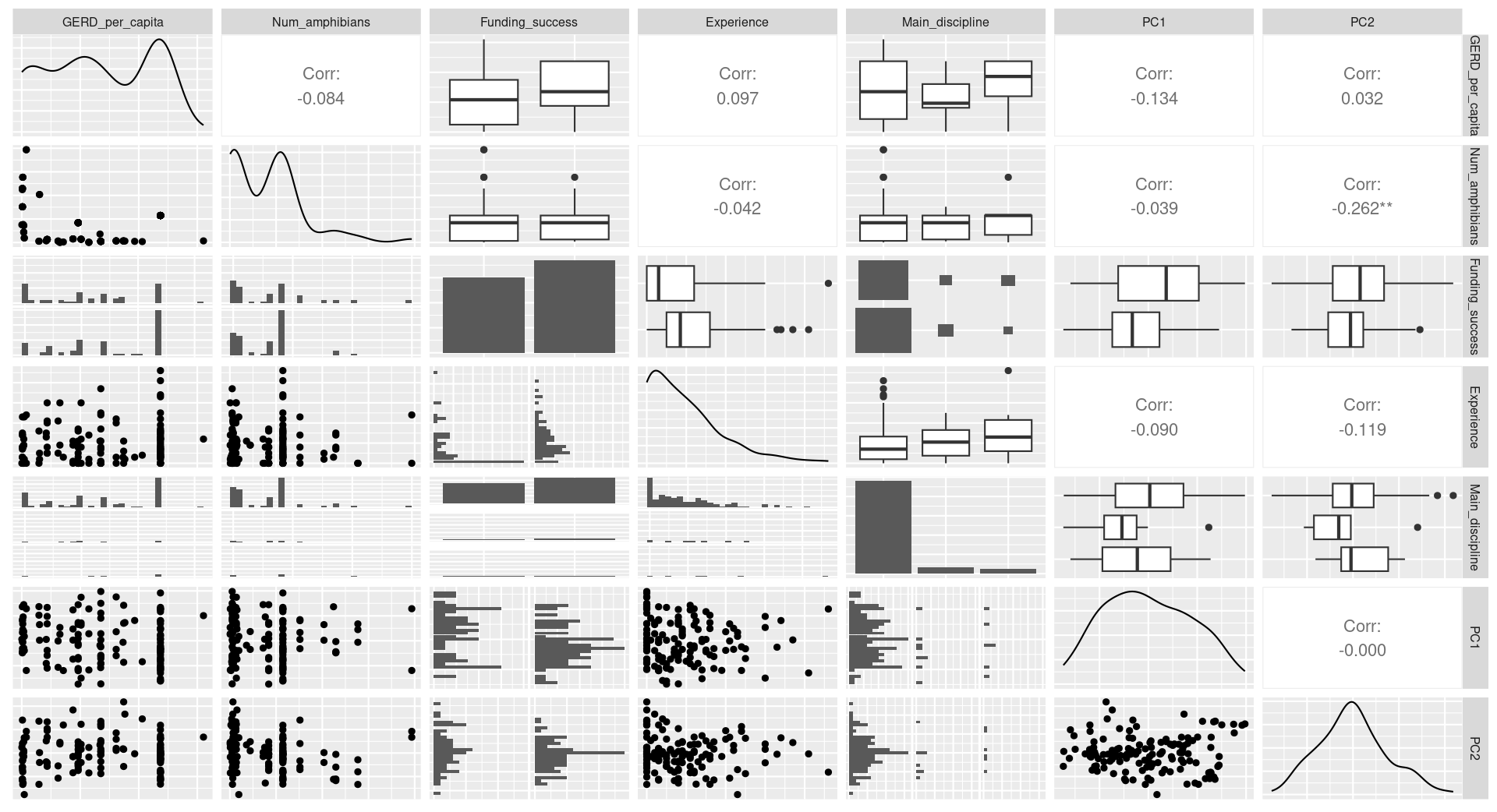

**Figure S3. Relations and distributions of the variables to model responses of the Amphibian Genomics Consortium (AGC) survey.** Orthogonal dimensions or principal components (PC) as independent variables are compared to the explanatory variables. Information about the number of amphibians per country was obtained from the AmphibiaWeb. The research and development expenditure of the gross domestic product (GERD) per capita was calculated using the information from the UNESCO and World bank websites provided about the most recent year for each country.

**References**

1.	Gregory TR: **Animal genome size database**. 2024.

2.	Liedtke HC, Gower DJ, Wilkinson M, Gomez-Mestre I: **Macroevolutionary shift in the size of amphibian genomes and the role of life history and climate**. *Nature Ecology & Evolution* 2018, **2**(11):1792-1799.

3.	Jetz W, Pyron RA: **The interplay of past diversification and evolutionary isolation with present imperilment across the amphibian tree of life**. *Nature Ecology & Evolution* 2018, **2**(5):850-858.

4.	Challis R, Kumar S, Sotero-Caio C, Brown M, Blaxter M: **Genomes on a Tree (GoaT): A versatile, scalable search engine for genomic and sequencing project metadata across the eukaryotic tree of life [version 1; peer review: 2 approved]**. *Wellcome Open Research* 2023, **8**(24).

5.	Revell LJ: **phytools 2.0: an updated R ecosystem for phylogenetic comparative methods (and other things)**. *PeerJ* 2024, **12**:e16505.

6.	Kassambara A: **Factoextra: extract and visualize the results of multivariate data analyses**. *R package version* 2016, **1**.
